# Measuring and reducing segmentation-driven cell-type labeling errors in imaging spatial transcriptomics

**DOI:** 10.64898/2026.11.23.768643

**Authors:** Vlad Elgart, Joseph Loscalzo

## Abstract

Imaging spatial transcriptomics labels each segmented cell from transcripts that include some of its neighbors’ RNA. We labeled each cell from its nucleus and its whole mask at equal depth, with two nuclear halves as a null. Whole-cell counts drew small, RNA-poor cells toward their larger neighbors’ labels. On three platforms, relative to nuclear labels at equal depth, T and NK cells lost 23–53% of their labels in six datasets of five tissues and 4–9% in renal cell carcinoma; this label-only measure matched same-cell protein markers in five donors and, where it differed, understated the loss. In gut, labels alone placed the loss at 58% of T cells among epithelial cells, compared with 24% in the lamina propria. The error lowered immune contact counts mainly by removing T cells; random relabeling, a common robustness test, raised them. Nuclear labels, where nuclei held enough counts, recovered many lost T cells. Imaged nuclei also carried some neighbor RNA and isolated nuclei carried none, indicating that these losses are lower bounds.

## Introduction

Cell-type labels are the common currency of imaging spatial transcriptomics: composition, colocalization, immune infiltration, and cellular neighborhoods are all computed from them [1]. Each label is called from the transcripts assigned to a segmented cell, and no segmentation assigns every transcript to the correct cell [2–4]. A T cell wedged between epithelial cells can receive a mask that includes some of their cytoplasm, and its label is then called from both. Methods that correct this admixture clean the expression and mostly keep the labels [3, 4], even though the labels were called from the same admixed counts; SPLIT changes a label only to swap a cell’s primary and secondary types where cells of similar expression carry the secondary type [4]. resolVI relabels cells, using a classifier trained on the existing labels, and shows the cells whose label then differs from the original annotation [5]. Jones et al. reported that small, RNA-poor cells such as T cells and neutrophils take the labels of larger neighbors, and that nuclear segmentation showed more T cells than expanded masks [6]. These corrections, and new segmentations, change the counts or the labels, but, as far as we know, none separates the labels that the admixture changed from those that counting noise changed in a given dataset, or checks them against an independent measurement on the same cells. We provide such a measure. It uses each cell’s own nuclear counts, needs no protein markers or spatial annotation, only a single-cell reference, and agreed with protein markers on the same cells.

How much this matters for the labels, and for the analyses built on them, has not been determined. Robustness is often tested by perturbing a random share of labels, wherever the cells sit [7–9] (Supplementary Table S1). By analogy, epidemiological studies distinguish two kinds of misclassification [10–12]: non-differential error does not depend on the exposure and mostly weakens associations; differential error depends on it and can bias them either way. An error caused by a neighbor’s transcripts depends on the neighbor, and the neighbor is what contact statistics measure. Nuclear counts are usually taken to define cell populations, with cytoplasmic counts expected to sharpen clustering and labeling; the default expansion of Xenium masks follows this reasoning [13]. On Xenium, expanded masks regrouped astrocytes, microglia, and oligodendrocytes by brain region rather than by type, which the authors attributed to domain-specific expression [13]. Segmentation benchmarks score methods against a single-cell reference, protein, or manual annotation [14], but most datasets have no protein markers or annotation, and manual annotations carry their own biases [15]. Thus, users have had no way to estimate the error in their own data.

Xenium and CosMx record whether each transcript lies over the nucleus, which allows a comparison inside every cell. Neighbors’ RNA reaches the nucleus less often than the rest of the mask, and, therefore, labels from nuclear and whole-cell counts at equal depth should agree unless the rest of the mask carries foreign RNA. We built this comparison into a test with a null for counting noise, and applied it to seven tissues on three platforms, three of them with protein markers measured on the same cells.

We find that whole-cell counts pull labels toward the contacting neighbor, and that the labels alone measure this error as a lower bound: the label-based measure agreed with protein markers across segmentations, tissues, and cell classes and, where it differed, understated the loss. How much a new segmentation helps depends on the masks it replaces. Small cells that border larger ones lose their labels, most of all where they sit inside a layer of larger cells: in gut, whole-cell labels lost most of the T cells within the epithelial layer and a quarter of those in the lamina propria. Studies that count lymphocytes within the epithelial layer or tumor-infiltrating T cells measure the cells most exposed to the error. Contact counts fall with the lost cells, the opposite of what random relabeling does. Labeling small cells from their nuclear counts where the nucleus holds enough of them, and from the whole cell otherwise, recovers T cells where whole-cell counts lose the most.

## Results

### Whole-cell counts draw labels toward the neighbor

To separate errors caused by the mask from counting noise, we labeled each cell three times from its own transcripts (Fig. 1a; Methods). Two random halves of the nuclear transcripts, A and B, differ only by chance, and their labels define the null. The third label comes from the rest of the mask, thinned to the depth of B. If the rest of the mask added only noise, this label would differ from that of A about as often as the label of B does. For each change beyond the null, we asked whether the new label was that of a contacting neighbor. The extra-change ratio compares how often this happened with how often it would happen if changes ignored the neighbors; it is one when the neighbors play no part.

**Figure 1.**
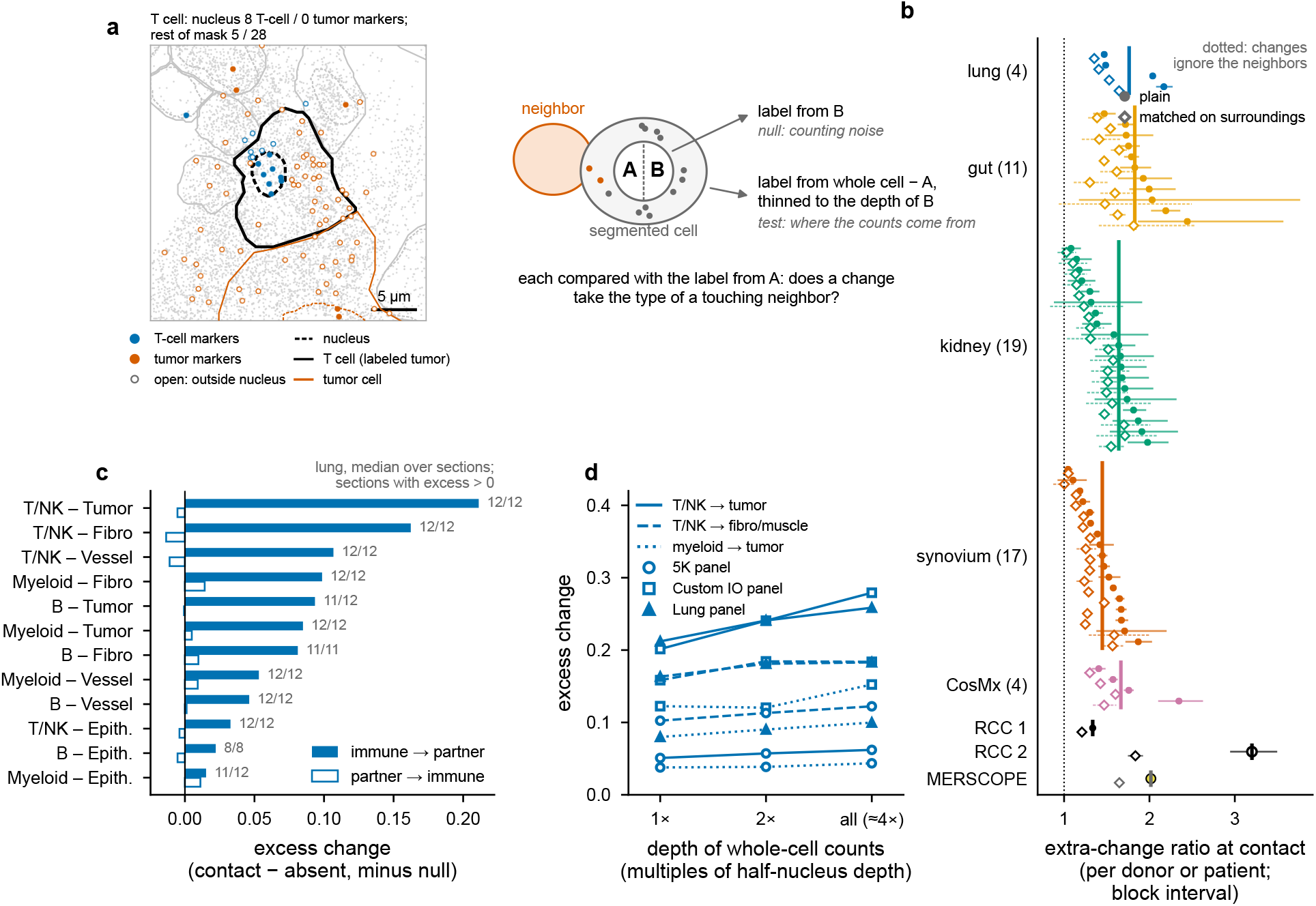
Whole-cell counts draw labels toward the neighboring cell. a, Left, a T cell in lung cancer (black outline) next to a tumor cell (vermilion outline); filled points, transcripts over a nucleus (dashed); open points, transcripts elsewhere; blue, T-cell markers; vermilion, tumor markers. Its nuclear counts are labeled T/NK and its whole-cell counts tumor (Supplementary Methods). Right, each cell is labeled from two halves of its nuclear counts (A, B) and from its whole-cell counts minus A, thinned to the depth of B; B against A is the counting-noise null. b, Extra-change ratio at contact: the observed share of label changes beyond the null that take a contacting neighbor’s type over the share expected if changes ignored the neighbors; per evaluable donor or patient with 95% block-bootstrap intervals; bars, medians; dotted line, one; open diamonds, the same ratio with the expected share matched on the composition of cells 30–100 µm away and the number of contacts. RCC 1, 2, the two renal cell carcinoma sections; MERSCOPE, the uterine cancer section. c, Excess change from an immune type toward a contacting non-immune partner (filled) and in the reverse direction (open): the rate at contact minus the rate without, minus the same contrast in the null; lung, medians over sections, with the number of sections in which the excess is positive. d, Excess change against the depth of the whole-cell counts (multiples of the half-nucleus depth), lung, per panel; the 5K sections, segmented with a membrane stain, show the smaller excess.

Label changes went to the neighbor more often than chance (Fig. 1b). The ratio was above one in all 51 evaluable donors and patients of four Xenium tissues: lung cancer [4], gut mucosa [16], kidney [17], and rheumatoid synovium [18]. It was also above one in lung cancer on CosMx [19], in two renal cell carcinoma sections with protein markers on the same cells [20, 21], and in a uterine cancer section on MERSCOPE [22]. Because cell types cluster in tissue, a change toward a neighbor’s type could reflect the surrounding region rather than the contact, for example, if expanded masks capture regional expression [13]. Matching each change on the composition of cells 30–100 µm away left most of the excess (Fig. 1b, open diamonds; Methods), so label changes follow the contacting cells. The result held with a second labeler, without uncertain labels, with contact defined by shared cell boundaries, and on labels confirmed by two further thirds of the nucleus (Supplementary Table S2). For synovium, we defined expected ranges from the other tissues before labeling: outcomes that follow from the measurement held, and some that depend on tissue architecture did not (Supplementary Table S3). A label audit showed that an over-detected probe had misled our first labeling of synovium (Supplementary Methods).

In the samples we examined, the pull ran mostly in one direction (Fig. 1c). In lung, a T or NK cell in contact with a tumor cell was labeled tumor in 0.26 of cases, compared with 0.04 in the null, whereas tumor cells in contact with T cells were not drawn toward T. More counts did not help: with all whole-cell counts, about four times the depth of the null, the excess did not shrink (Fig. 1d). The error, therefore, lies in what the mask contains rather than in a shortage of counts.

The test needs labels that are stable at nuclear depth. The two nuclear halves agreed on 0.56–0.99 of labels in the Xenium sections; in most pieces of a CosMx liver dataset, they agreed too rarely for the test to be read (Supplementary Note 2).

### The labels alone track the loss that protein markers measure

We checked the measure against protein expression on the same cells, which we had in three forms: immunohistochemistry for CD8 on two lung sections after the Xenium run, registered to the Xenium nuclei [4]; the 27-protein Xenium Protein assay on two renal cell carcinoma sections; and CD45RO imaged with the transcripts on a MERSCOPE section of uterine cancer. The markers identified individual cells rather than regions (Supplementary Methods). For each section and segmentation, we computed two ratios, both relative to nuclear half B at equal depth (Methods). The kept ratio uses labels only: the share of half A T/NK labels that whole-cell counts keep over the share that half B keeps. The protein marker recall ratio uses protein markers: the share of protein-positive T cells that whole-cell labels call T over the same share for half B.

Across nine segmentations of two lung sections, from nuclei expanded by 1 to 15 µm to the deposited masks and Proseg, the two ratios fell and leveled off together (Fig. 2a), although the labels group T and NK cells and the protein marker identifies only CD8^+^ cells. Over all 30 comparisons, they differed on average by at most 0.02 per donor (Fig. 2b), and correcting the lung recalls for chance matches barely changed them (Supplementary Table S5). On MERSCOPE, with nuclei segmented from the DAPI image, the kept ratio was 0.51 and the protein marker recall ratio 0.50, a loss as large as the largest on Xenium.

**Figure 2.**
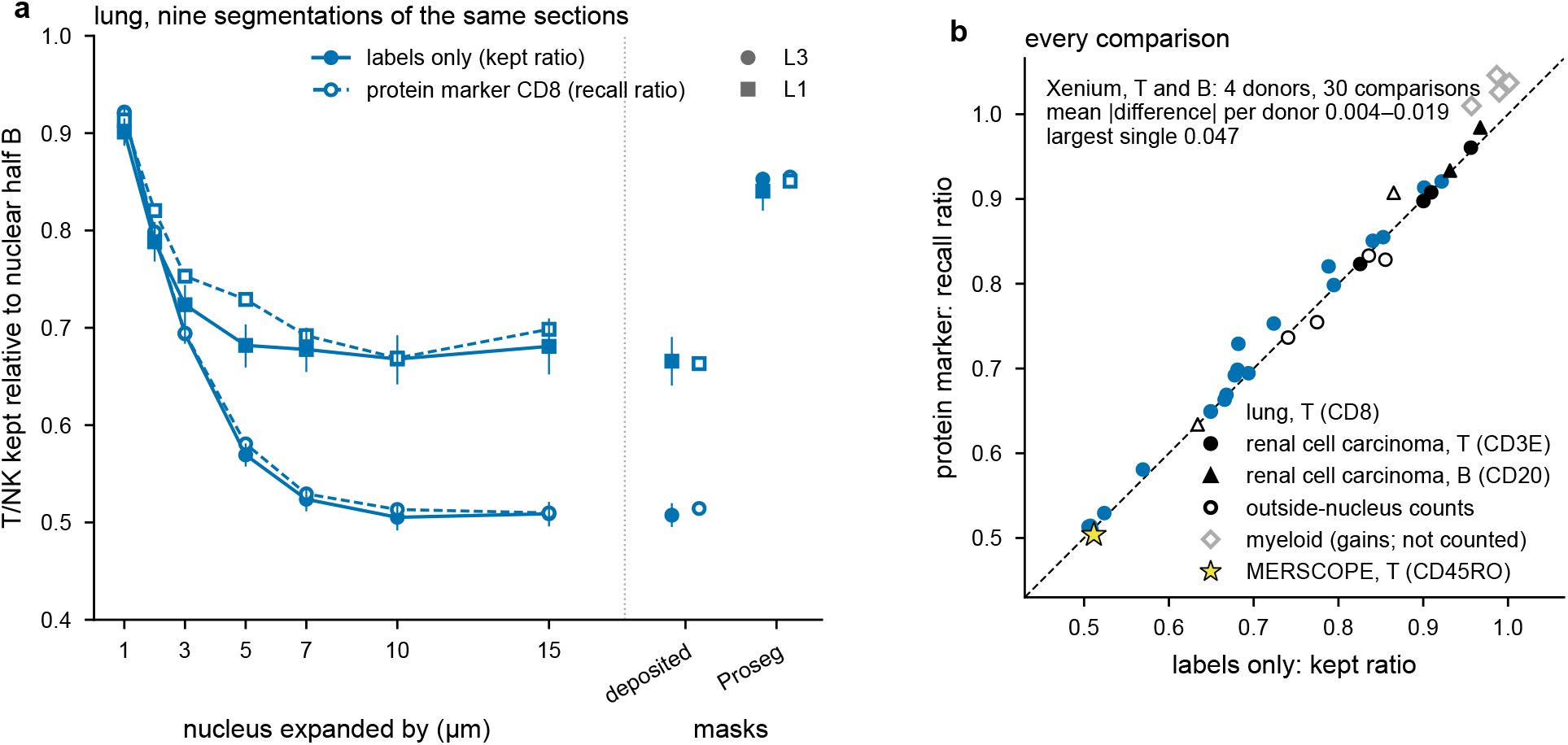
The labels alone track the loss that protein markers measure. Both measures relative to nuclear half B at equal depth. a, Lung sections L1 (squares) and L3 (circles), nine segmentations of the same cells: T/NK kept ratio from the labels (filled, solid; block intervals) and recall ratio of protein marker CD8^+^ cells (open, dashed), for nuclei expanded by 1–15 µm, the deposited masks, and Proseg. b, Every comparison: protein marker recall ratio against kept ratio, for T cells in lung (CD8, blue) and T and B cells in the two renal cell carcinoma sections (CD3E, CD20; black; deposited masks and, for T cells, Proseg), from whole-cell (filled) and outside-nucleus counts (open); gray, myeloid cells, which gain labels that the kept ratio does not count; yellow star, uterine cancer on MERSCOPE (T cells by CD45RO, whole-cell counts; block interval of the kept ratio); dashed, identity; inset, the range over the Xenium donors of the mean absolute difference of their T and B comparisons, and the largest single difference.

Part of this agreement is built in because the two ratios share most of their cells: in renal cell carcinoma, 0.63 and 0.62 of the T cells defined by protein markers carried a half A T/NK label (Supplementary Table S4). The comparison that shares no cells with the kept ratio is the recall of the T cells that half A called another type. In renal cell carcinoma, these cells kept their T label about as often as the kept ratio implies (0.87 and 0.94). In lung, they lost it more often (0.54 and 0.28, against kept ratios of 0.67 and 0.51): the T cells whose nuclei carry the weakest T-cell signal were lost most often, and the kept ratio, which cannot see them, understates the loss there. Protein markers also showed that the lost labels belonged to true T cells: lost CD8^+^ cells took on the types of the cells around them (Supplementary Table S6), and the share of true T cells among the T labels did not fall (Supplementary Table S7). The kept ratio counts labels lost, not labels gained, so it misses the gain of the myeloid class that absorbs them (Fig. 2b, gray).

The nuclear reference is not free of neighbor RNA. T/NK cells in contact with a partner cell carried more of the partner’s marker transcripts than T/NK cells away from one, in whole-cell counts and, to about half the extent, in nuclear counts, in all four Xenium tissues (Supplementary Fig. S1a; Supplementary Table S8). Mislabeled cells did not explain the nuclear excess, which remained with confident labels only and with the load counted in the nuclear half that the labels did not see. In renal cell carcinoma, with T cells and tumor cells typed by protein markers and no label, the nuclei held most of the whole cell’s excess. Nuclei dissociated and placed by Slide-GoTags [23] held none, and neither positional error larger than that of the method [24] nor the removal of cells explained the difference (Supplementary Fig. S1b). An imaged nucleus is a two-dimensional outline that also collects RNA above, below, and beside it [25], and, thus, the losses measured against nuclear labels are lower bounds.

### Segmentation sets the size of the error

If the mask causes the error, changing only the mask should change the error. We rebuilt the cells of the lung and kidney sections by expanding their nuclei by 1 to 15 µm, which leaves the nuclear counts, and hence the null, unchanged (Methods). One micrometer of expansion already removed 0.10–0.18 of T/NK labels, because half the gap to the nearest nucleus is about a micrometer or less (Methods). The loss leveled off from 5–7 µm (Fig. 3a), as did the loss measured by protein markers (Fig. 2a). Each outside transcript goes to the nearest nucleus, so expanded nuclei soon meet their neighbors’ expansions: half the gap to the sixth-nearest nucleus was 3.6–6.0 µm, and by 5 µm the masks held nearly all the outside counts that 15 µm gives (Methods). The deposited masks of the Lung and Custom IO panels are drawn by nucleus expansion, and they lost about as many labels as nuclei expanded by 15 µm.

**Figure 3.**
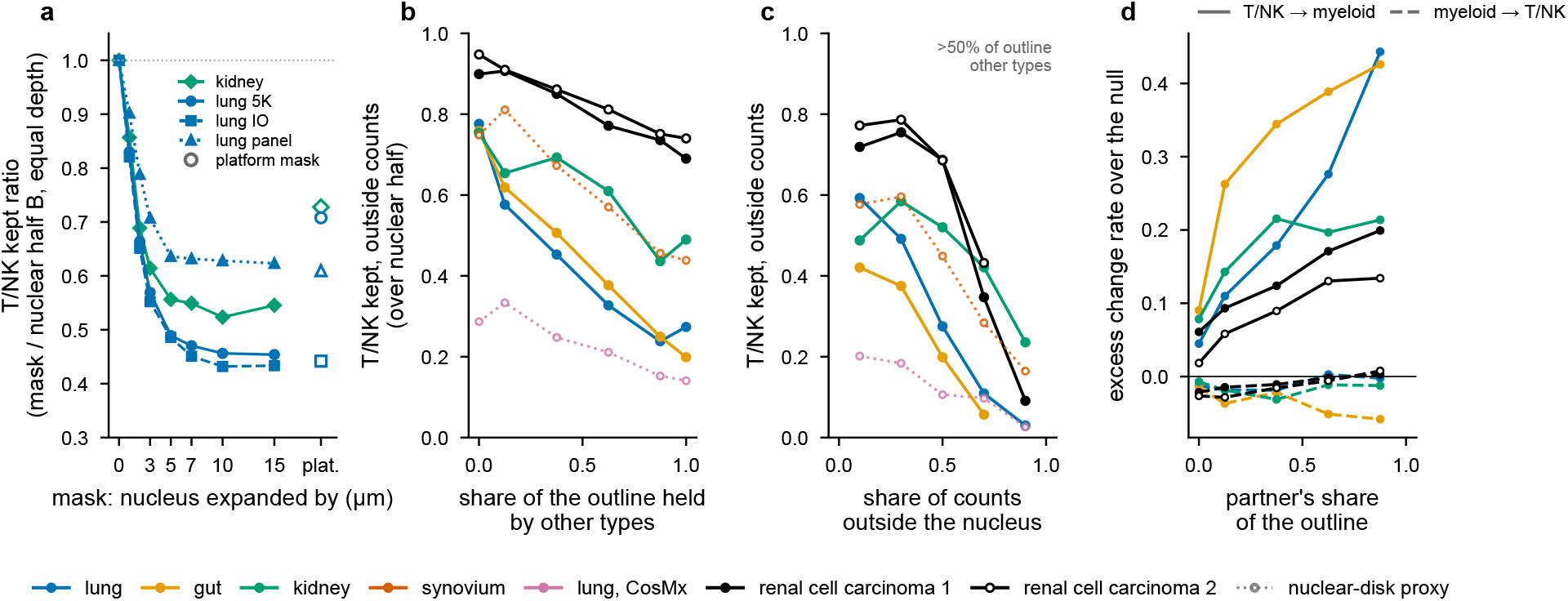
The mask collects the neighbors’ RNA. a, T/NK kept ratio of each mask relative to nuclear half B at equal depth, for nuclei expanded by 0–15 µm and for the platform’s masks (open; stain segmentation in lung 5K and kidney); medians over sections with at least 100 T/NK cells in the null (12 lung sections by panel, 9 kidney regions). b–d, Counts outside the nucleus only, medians over donors or patients (one section per renal cell carcinoma donor; donor 2 open): b, T/NK kept against the share of the outline held by other types, from the segmentation polygons (solid) or, for synovium and CosMx, which have none, from the nuclear-disk proxy (dotted, faint); c, against the share of counts outside the nucleus, for cells with more than half of their outline held by other types; d, polygon datasets, excess change T/NK *→* myeloid (solid) and myeloid *→* T/NK (dashed) against the partner’s share of the outline.

We then compared the segmentations that we could test on the same tissue (Table 1). In lung 5K, masks drawn from a membrane stain lost fewer labels than expanded nuclei (Fig. 3a); in the one gut patient imaged with and without the stain, they lost slightly fewer, within overlapping intervals. Two transcript-based methods, Proseg [6] and Baysor [2], cut the loss by about half to three quarters in the two lung sections, whose deposited masks are expanded nuclei, but recalled no more T cells in the two renal cell carcinoma sections, whose masks come from a segmentation stain. There, Proseg’s kept ratio was even lower, because Proseg also removes nuclear transcripts that it assigns to other cells, which makes the nuclear reference cleaner. With every segmentation, the remaining changes still went to the contacting neighbors (Supplementary Table S2), and on Baysor cells, too, the kept ratio and the protein marker recall ratio agreed.

**Table 1.**
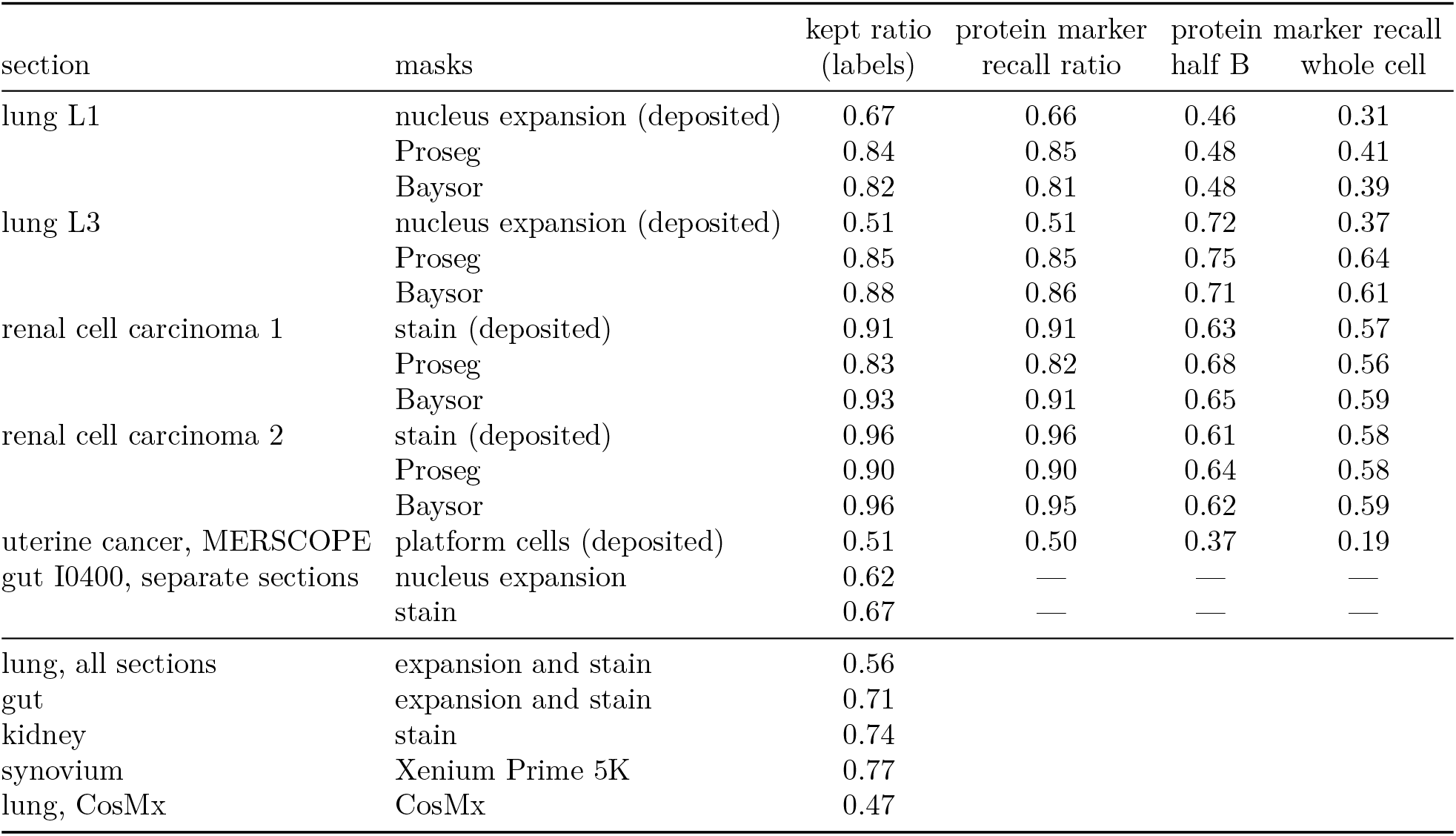
How many T/NK labels whole-cell counts lose, by cell segmentation. Ratios relative to nuclear half B at equal depth; one is no loss. Kept ratio, from labels only; protein marker recall ratio, the recall of protein-marker-positive cells (lung, CD8^+^ by immunohistochemistry; renal cell carcinoma, CD3E^+^; uterine cancer, CD45RO^+^ imaged in the same run) by whole-cell labels over that by half B; absolute recalls of half B and of whole-cell counts at equal depth. Uterine cancer: the platform’s cells, all of them. Lung and renal cell carcinoma rows compare segmentations on the cells both keep; the renal cell carcinoma deposited rows are rescored on the cells Proseg keeps. Baysor rows: the cells Baysor keeps; deposited masks on those cells kept 0.66 (L1), 0.51 (L3), 0.91 (renal cell carcinoma 1), and 0.96 (2). Gut I0400: separate sections of one patient, the only one with at least 100 T/NK cells in the null in both. Lower block: medians over donors or patients, deposited masks (Fig. 4a). Block intervals in Supplementary Table S2, unit counts in Supplementary Table S9, and protein cell counts in Supplementary Table S5.

|  |  | kept ratio | protein marker | protein marker recall |  |
|---|---|---|---|---|---|
| section | masks | (labels) | recall ratio | half B | whole cell |
| lung L1 | nucleus expansion (deposited) | 0.67 | 0.66 | 0.46 | 0.31 |
|  | Proseg | 0.84 | 0.85 | 0.48 | 0.41 |
|  | Baysor | 0.82 | 0.81 | 0.48 | 0.39 |
| lung L3 | nucleus expansion (deposited) | 0.51 | 0.51 | 0.72 | 0.37 |
|  | Proseg | 0.85 | 0.85 | 0.75 | 0.64 |
|  | Baysor | 0.88 | 0.86 | 0.71 | 0.61 |
| renal cell carcinoma 1 | stain (deposited) | 0.91 | 0.91 | 0.63 | 0.57 |
|  | Proseg | 0.83 | 0.82 | 0.68 | 0.56 |
|  | Baysor | 0.93 | 0.91 | 0.65 | 0.59 |
| renal cell carcinoma 2 | stain (deposited) | 0.96 | 0.96 | 0.61 | 0.58 |
|  | Proseg | 0.90 | 0.90 | 0.64 | 0.58 |
|  | Baysor | 0.96 | 0.95 | 0.62 | 0.59 |
| uterine cancer, MERSCOPE | platform cells (deposited) | 0.51 | 0.50 | 0.37 | 0.19 |
| gut I0400, separate sections | nucleus expansion | 0.62 | — | — | — |
|  | stain | 0.67 | — | — | — |
| lung, all sections | expansion and stain | 0.56 |  |  |  |
| gut | expansion and stain | 0.71 |  |  |  |
| kidney | stain | 0.74 |  |  |  |
| synovium | Xenium Prime 5K | 0.77 |  |  |  |
| lung, CosMx | CosMx | 0.47 |  |  |  |

What a new method gains, therefore, depends on the masks it replaces; in a benchmark of 11 methods on 15 datasets, no method was best across datasets [14]. The kept ratio measures the gain on the data at hand. It ranks two segmentations fairly when they share the nuclear counts, as nuclear expansions do; a method that also changes the nucleus needs protein markers or another external reference.

### Small cells bordering larger ones lose their labels

Within each segmentation, the error was not distributed evenly over cells. T/NK labels were kept almost as often as in the null when a cell’s outline made contact only with cells of its own type, and less often as other types held more of it, in every dataset with segmentation polygons (Fig. 3b). A larger share of the mask outside the nucleus added to the loss (Fig. 3c). Protein markers confirmed the gradient without relying on any label: in both renal cell carcinoma donors, the loss of T cells defined by protein markers rose steadily with the share of their outline held by CD45-negative cells, also within bins of the composition of the surrounding region (Fig. 4c; Supplementary Table S10). At the same exposure, the two donors lost the same share; donor 2 lost less overall because more of its T cells were in contact only with immune cells. Donor 1 also lost T cells that had no non-immune contact, a loss that the outline does not explain.

**Figure 4.**
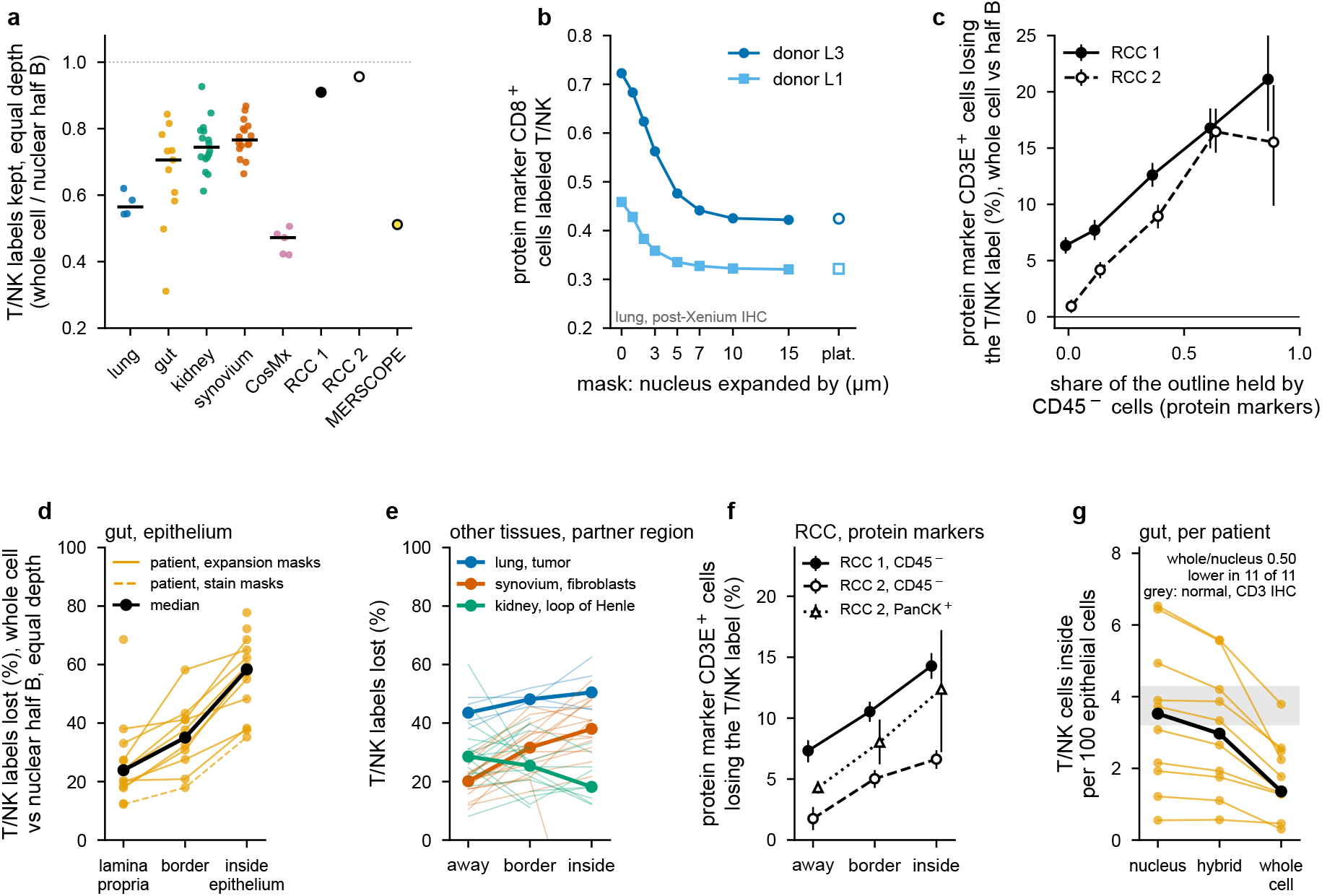
T cells are undercounted when they border other cells, most among epithelial cells. a, T/NK kept ratio of whole-cell counts relative to nuclear half B, at equal depth, per donor or patient with at least 100 T/NK cells kept in the null (bars, medians) and for the single sections of renal cell carcinoma (RCC 1 and 2) and uterine cancer (MERSCOPE), with block intervals. b, Protein marker CD8^+^ cells (post-Xenium immunohistochemistry, lung donors L1 and L3) labeled T/NK, by mask; open, platform masks. c, Renal cell carcinoma, both donors (donor 2 open, dashed): protein marker CD3E^+^ cells losing the T/NK label, whole-cell counts against nuclear half B, by the share of the cell’s outline (segmentation polygon) held by CD45*^−^* cells, with 95% block intervals; bins of at least 500 cells shown (the slopes in Supplementary Table S10 use bins of at least 200). d, Gut: T/NK labels lost (whole-cell counts against nuclear half B, equal depth) by where the cell sits, from the share of epithelial cells among the labeled cells within 25 µm: lamina propria (< 0.2), border (0.2–0.5), and inside, among epithelial cells (*≥* 0.5); one line per patient (solid, nucleus-expansion masks; dashed, stain masks), medians in black. e, The same for lung (tumor), synovium (fibroblasts), and kidney (loop of Henle); thin lines, donors or patients; thick, medians; few kidney T/NK cells sit inside the tubules (Supplementary Table S11). f, Renal cell carcinoma, protein truth: protein marker CD3E^+^ cells losing the T/NK label, by region cells typed by protein (CD45*^−^*, both donors; PanCK^+^, donor 2), with 95% tile-bootstrap intervals. g, Gut: T/NK cells among epithelial cells (inside, as in d), per 100 epithelial cells, per patient, from nuclear, hybrid, and whole-cell labels; medians in black; grey, the interquartile range of counts by image analysis of CD3-stained surface epithelium in normal colon [26]; ours include crypt epithelium.

The exchange was asymmetric. T/NK cells took the myeloid label more often the more of their outline myeloid cells held, but myeloid cells did not take the T/NK label at any share (Fig. 3d). This pattern is what one would expect from the mask: a T cell has few transcripts of its own, so the RNA of a large, RNA-rich neighbor can outweigh them, whereas a few foreign transcripts barely change a cell that has many. Marker counts agreed. In each of the five datasets, the cytoplasm of T/NK cells in contact with other types carried more of their marker counts, above the baseline set by the nucleus, than the cytoplasm of T/NK cells in contact only with T/NK cells (Methods). Even these unexposed T cells carried some excess, which points to foreign RNA from beyond the contacting cells, such as cells above or below the section plane [25]. This foreign load, measured on one half of the whole-cell counts, predicted the label changes made from the other half (Supplementary Fig. S2).

The loss did not depend on the classifier’s settings (Supplementary Table S2); its size followed tissue architecture. Whole-cell labels kept roughly half to three quarters of the T/NK labels across six datasets on three platforms (Table 1; Fig. 4a), and sections differed mainly in how many T cells were exposed rather than in how much an exposed T cell lost. The loss was not specific to T cells. B and mast cells lost labels in every tissue where they could be measured, and in kidney the thin mural cells wrapped around vessels lost the most (kept ratio 0.64, against 0.76 for T/NK cells in the same computation), whereas tumor, tubular, and gut epithelial labels were kept (Supplementary Table S20). Labels made with protein as well showed the same loss: the vendor’s labels for renal cell carcinoma donor 2 lost the most exposed T cells as often as our whole-cell labels did (Supplementary Fig. S3).

### T cells among epithelial cells lose the most labels

The outline describes a cell’s immediate border. To ask where in the tissue the loss falls, we placed each T/NK cell by the share of partner cells among the labeled cells within 25 µm and called it away from, at the border of, or inside the partner region (Methods), a grouping by region rather than by contact. In gut, with epithelium as the partner, whole-cell labels at equal depth lost 24% of T/NK labels in the lamina propria, 35% at the border of the epithelium, and 58% inside it, within the epithelial layer (Fig. 4d). The loss was higher inside than away in 11 of 11 patients, with nucleus-expansion masks and with masks from a segmentation stain alike (Supplementary Table S11): the masks set the level of the loss, and the tissue set where it fell. In synovium, T/NK cells surrounded by fibroblasts likewise lost more labels than those elsewhere (15 of 16 patients), and in lung, T/NK cells inside tumor nests lost somewhat more (3 of 4 donors; Fig. 4e). In kidney, few T/NK cells sat inside the tubules, and they lost no more labels than those away; counted against all non-immune cells, those surrounded by them lost more (Supplementary Tables S11 and S12).

A T/NK call among epithelial cells could instead be an epithelial nucleus carrying T-cell RNA, for example, from a cell above or below the section plane, and whole-cell counts would then correct its label rather than lose it. Most inside calls did not look like epithelial nuclei (Supplementary Table S13). Their nuclei were smaller than epithelial nuclei, held T-cell rather than epithelial marker counts, and carried cytotoxic transcripts, as lymphocytes within the epithelial layer do; nuclear half B, which shares no transcript with half A, called them T/NK at least as often as it called the T/NK cells of the lamina propria. Some may still be epithelial: the runner-up label of half A was epithelial for 0.22 of them. Their whole-cell masks, in contrast, were larger than those of T/NK cells in the lamina propria and about as large as an epithelial cell’s: the mask of a T cell wedged between epithelial cells reaches into their cytoplasm. T/NK calls inside lung tumor nests were likewise T-sized and cytotoxic.

Protein markers test the identity without any label. In the two renal cell carcinomas, with T cells called by CD3E protein and region cells typed by protein as CD45-negative, whole-cell labels lost more T cells inside non-immune regions than away from them, and most inside pan-cytokeratin-positive tumor (Fig. 4f; Supplementary Table S11). The gradient was shallower than in gut, as was the loss overall in these sections (Table 1), but the loss fell in the same place.

Whether whole-cell labels lose a T cell thus depends on where it sits, and in gut the cells they lose are cytotoxic T cells among epithelial cells. In the units pathologists use for lymphocytes within the epithelial layer, per 100 epithelial cells, whole-cell labels gave about half as many T cells among epithelial cells as nuclear labels (median ratio 0.50; lower in 11 of 11 patients; Fig. 4g): a median of 1.4 per 100, against 3.5 with nuclear labels, close to the about four that CD3 staining finds in normal ileum and colon [26, 27]. Hybrid labels came close to nuclear labels (median ratio 0.89). Our counts include crypt epithelium, come from IBD tissue, and are made in two-dimensional sections, so the agreement in level is approximate (Supplementary Methods). Within a region, a cell’s state mattered little: after matching cells on nuclear counts, no gene set changed the loss in the same direction in every tissue, whereas the contrast between inside and away remained in gut and synovium and was smaller in lung and kidney (Supplementary Methods; Supplementary Table S12). B cells showed the same gradient, but their calls near partner cells were uncertain (Supplementary Table S11).

### Contact counts fall with the lost T cells; random relabeling moves them in the opposite direction

Contact statistics follow the cells that are lost. With whole-cell labels at the depth of nuclear half B, T/NK contacts per 100 partner cells fell by about half on lung tumor and on gut epithelium, and they fell in kidney and synovium, as well (Fig. 5a). To ask how much of this fall comes from where the lost labels sit, we placed the same label changes at random, with the real confusion rates and every partner cell kept at its half B label (Methods). The real fall exceeded this control in 50 of 56 donors and patients. The fall, therefore, has two parts: fewer T/NK cells, which accounts for most of it (Fig. 5a, hatched bars), and lost labels placed at contact more often than chance, which accounts for the gap between the filled and hatched bars. Statistics normalized to the immune class, such as the share of T cells in contact with the partner, changed little where the loss was spread over the tissue, whereas raw counts fell with the class. Gut is the exception: there the loss falls on the T cells among epithelial cells, so even the share of T cells in contact with epithelium is biased. When the partners were relabeled at random, as well, as a label-only control does by default, the same changes raised T/NK contacts in 18 of 56 donors and patients (Fig. 5a, light bars).

**Figure 5.**
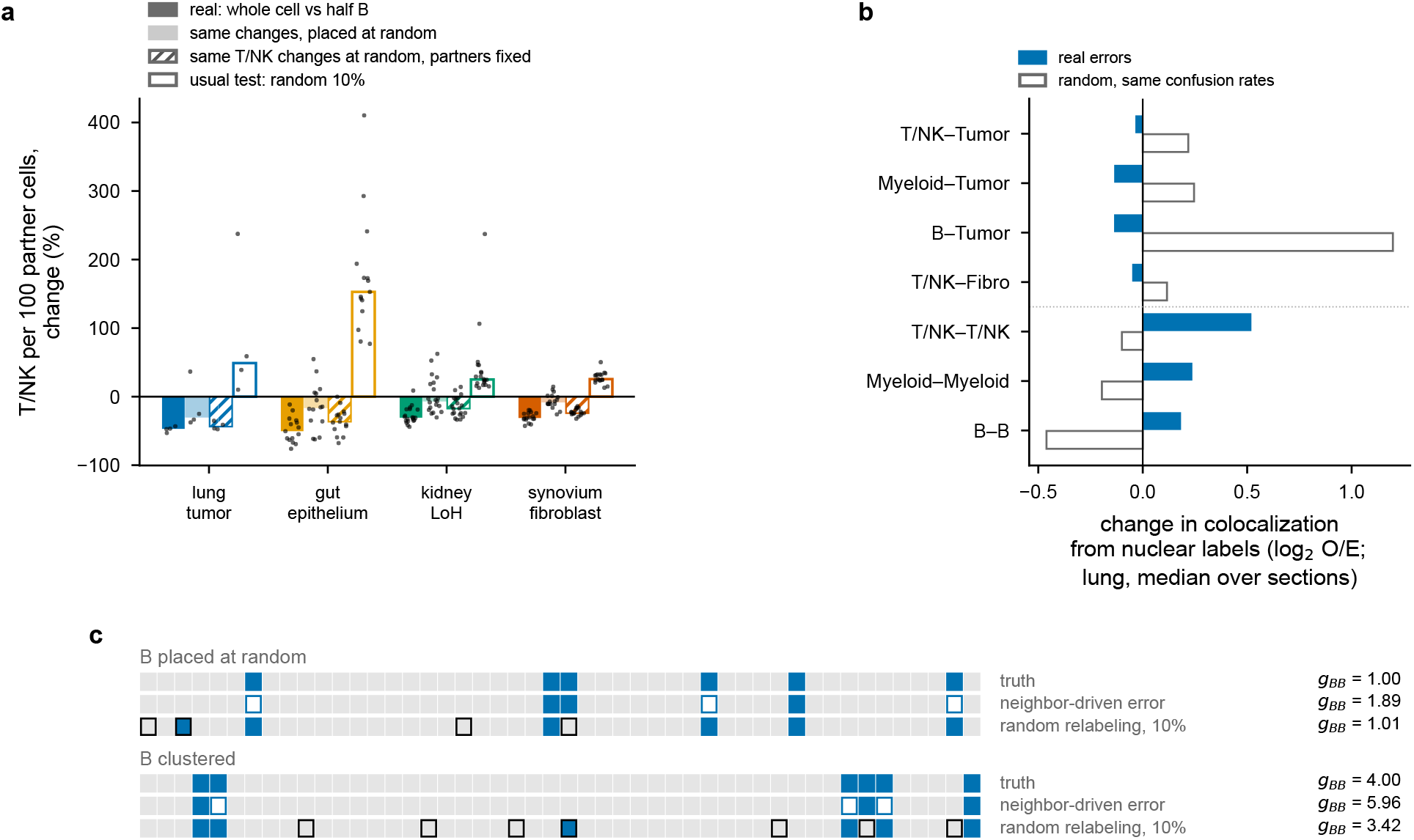
Real errors lower contact counts mainly by shrinking the T/NK class; random relabeling moves them in the opposite direction. a, T/NK cells contacting the partner per 100 partner cells: whole-cell labels at the depth of nuclear half B, change from the half B labels (filled); the half B labels relabeled at random with the real confusion rates, partner cells included (light) or kept at their half B labels (hatched), change from the half B labels; and the whole-cell labels after random relabeling of 10% of cells, change from the whole-cell labels (open); bars, medians over donors or patients; dots, donors or patients; LoH, loop of Henle. b, Change in colocalization from the nuclear labels, lung, medians over sections: whole-cell labels (filled) and random relabeling of the nuclear labels at the same confusion rates (open). c, A line of cells of two types (blue, B; gray, A) in which a B cell takes the label A with probability 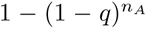, where *n_A_* is its number of A neighbors (*q* = 0.3), the simplest form of the contact dose in Fig. 3b. Shown on a stretch of 48 cells: the truth, after neighbor-driven error (open, B cells given the label A), and after random relabeling of 10% of cells (outlined), for B placed at random and clustered. Right, the B–B pair enrichment *g_BB_* over one million cells; with B placed at random, 0.53 of B labels survived the neighbor-driven error. Random relabeling preserves composition. The model places all error at contacts; in our data, the positional part was small where the loss was spread over the tissue and large where small cells sit inside a layer of larger ones (Results). The placement stress test is in Supplementary Fig. S4.

In the two renal cell carcinomas, protein markers allow the same checks without labels. With partners typed by protein markers, the positional part was about half of what the label-only check gives, so in the Xenium tissues that check likely overstates it; whole-cell counts lowered the contacts with CD45-negative cells but not the share of T cells in contact with them (Supplementary Table S14).

Random relabeling, a common robustness test, moved the same statistics in the opposite direction: reassigning a tenth of the labels at random raised T/NK contacts with the partner in every donor and patient (Fig. 5a, b). Placing the real number of errors on the cells with the most contacts reproduced the sign of the real change for more type pairs than random placement did (Supplementary Fig. S4). Random errors scatter immune labels into the partner compartment, whereas real errors remove them from it. A line of cells shows why (Fig. 5c). If a cell takes its neighbors’ label more often the more such neighbors it has, the cells that border the other type are lost first, and those that keep their label are the ones surrounded by their own type. Their aggregation then appears to rise, whether they were placed at random or clustered, whereas random relabeling leaves it unchanged or moves it toward independence, as non-differential error does [10, 12]. For a depleted pair, such as T cells and tumor, the two errors push in opposite directions, so a claim of immune exclusion that survives random relabeling has not yet faced the error that segmentation makes. In our data, no enrichment call changed sign; claims of depletion are the ones most at risk.

### What corrects the error, and at what cost

Users label from full counts, so the practical question is which counts recover the most cells at full depth. A nucleus labels its own cell well, but it needs enough counts to be labeled: only 0.79 and 0.71 of T cells defined by protein markers in renal cell carcinoma had a label from the whole nucleus. We, therefore, labeled each cell from its nucleus where the nucleus held enough counts, and from the whole cell otherwise. With unlabeled cells counted as missed, this hybrid recalled at least as many T cells defined by protein markers as either source alone in every section, at nearly the same precision (Supplementary Table S16). It gained most where whole-cell counts lost most: in lung, it recalled 0.39 and 0.66 of CD8^+^ cells, compared with 0.31 and 0.42 from the whole cell. Myeloid cells, which are large and RNA-rich, were the exception: whole-cell labels recalled more of them (Supplementary Table S16). In these data, small classes were best labeled from the nucleus, with the whole cell for cells whose nucleus held too few counts, and large classes from the whole cell.

At equal depth, too, the nucleus did better for the cells it labeled. In every tissue, half of a cell’s nuclear counts agreed with the other half more often than the same half with the cell’s outside counts added (Fig. 6b), and in both renal cell carcinoma donors, nuclear labels recalled more T, CD8, CD4, and B cells than any count set that included the cytoplasm (Fig. 6d; Supplementary Table S15).

**Figure 6.**
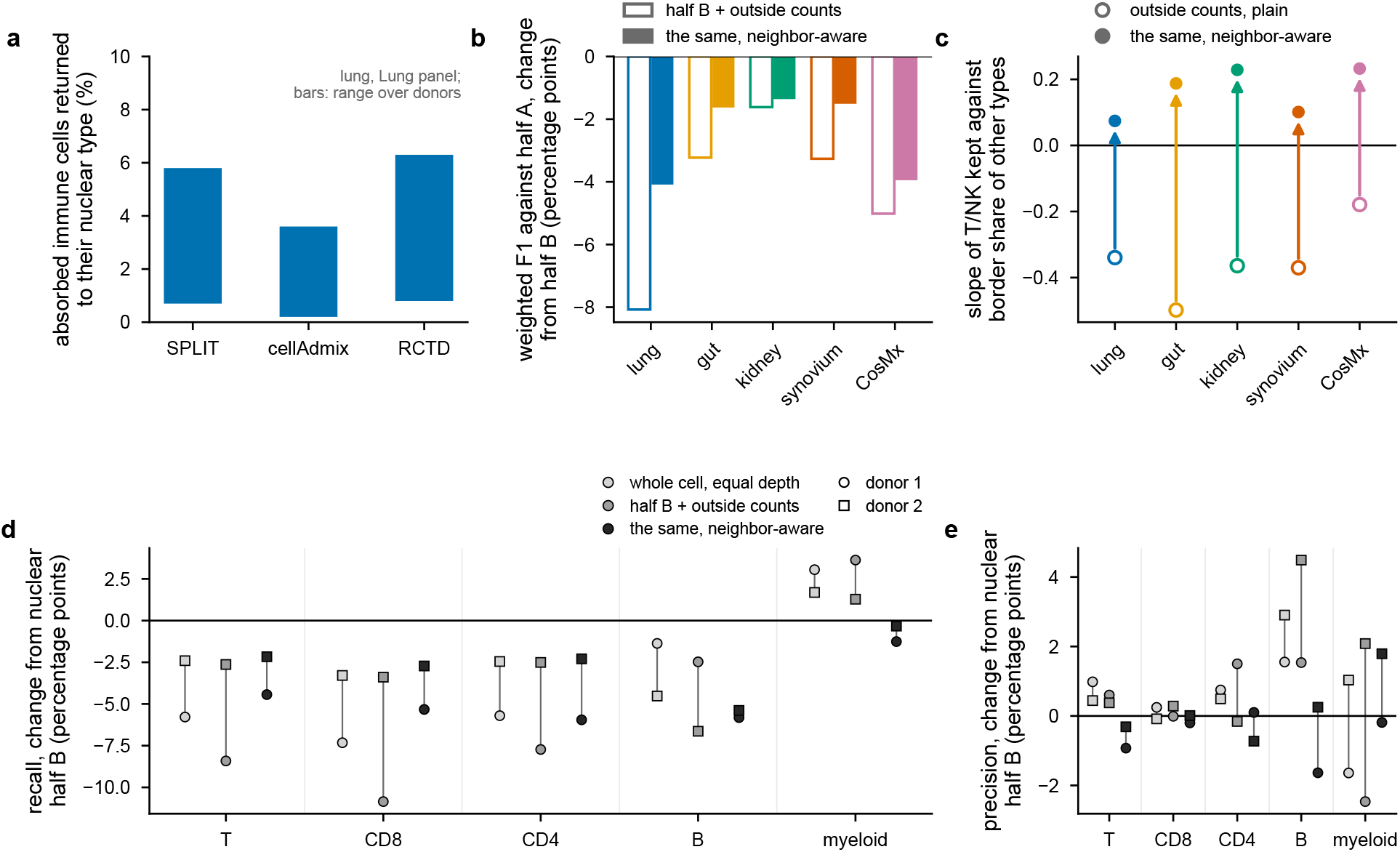
What corrects the error, and at what cost. a, Share of immune cells absorbed into a non-immune type that each correction returns to the nuclear type (lung, Lung panel; bars span the range over donors). b, Change in weighted F1 against nuclear half A from that of half B, for half B plus the outside counts (open) and the same with the neighbor-aware likelihood (filled). c, Slope of T/NK labels kept by outside counts against the share of the border held by other types (nuclear-disk proxy), plain (open) and neighbor-aware (filled): without the nucleus, the model reverses the dependence in every dataset (curves in Supplementary Fig. S5). d, e, Renal cell carcinoma, two donors (circles, donor 1; squares, donor 2; lines join the two), every class typed by protein on the same cells: change in recall (d) and precision (e) from nuclear half B for whole-cell counts at equal depth, half B plus the counts outside the nucleus, and the same counts with the neighbor-aware likelihood.

Published corrections did not close the gap. Most remove admixed counts given each cell’s label, such that they cannot return a cell that already carries an incorrect label; SPLIT-shift, the exception, only swaps a cell’s primary and secondary types. RCTD [28], SPLIT [4], and cellAdmix [3], run on five lung sections and followed by relabeling, returned only a few percent of absorbed immune cells to their nuclear type (Fig. 6a; Methods), and, by protein markers, SPLIT and RCTD lowered the share of CD8^+^ cells labeled T/NK (Supplementary Table S5). We then asked whether modeling the foreign RNA could make the cytoplasm usable. A likelihood that expects each cell’s outside counts to mix its own profile with the RNA of its neighbors, weighted by their share of the border, recovered some of the lost T cells in the two renal cell carcinomas, at a cost in the recall of other classes, and never reached the nucleus alone, even with its weights changed (Fig. 6d, e; Methods). Without the nucleus, it reversed the error instead of removing it (Fig. 6c). From the counts outside the nucleus alone, “a cell of type *a* holding RNA of type *b*” and “a cell of type *b* holding RNA of type *a*” explain the same counts; only a part of the cell its neighbors reach least, such as the nucleus, or an independent measurement such as a protein marker, separates them. resolVI [5], which models each cell’s counts as a mixture of its own expression, its neighbors’, and background and relabels every cell, recalled more T cells defined by protein markers than nuclear labels in every run and, on the windows of its first run, more than any of our labelings, even with our recall counted only over the cells our labels covered (0.72 and 0.73, against 0.67 and 0.65 for hybrid labels); it lowered T precision (0.90 and 0.87, against 0.92 and 0.92) and, depending on its filter and seed, the precision or recall of other classes (Methods; Supplementary Table S17). In our data, these models traded one error for another, so their net effect has to be measured on each dataset.

## Discussion

Segmented cells carry some of their neighbors’ RNA, and labels called from these counts are drawn toward the neighbors. By analogy with the language of epidemiology, the error is differential: it depends on the same exposure that contact statistics measure. Because it falls where small cells sit among larger ones, its size in a given study depends on tissue architecture as much as on the segmentation. The same signal can reach methods that read spatial context from a cell’s own expression. Nicheformer, for example, predicts the cell-type composition of a cell’s neighborhood from its counts [29]; in imaging data, part of that information can be neighbors’ RNA in the cell’s mask. Its benchmarks include the CosMx lung cancer data [19], in which whole-cell labels kept the fewest T/NK labels here (Table 1).

The most useful finding may be that this error can be measured in the data where it arises. The kept ratio needs only the nucleus flag that Xenium and CosMx record, or nuclei segmented from the DAPI image on MERSCOPE. It agreed with the loss that protein markers measured, across segmentations and for both classes that lose labels, and where the two differed, it understated the loss. With a reference classifier applied to the nuclear halves and to the whole cell, users can, therefore, estimate how many T cells their segmentation costs them relative to nuclear labels, as a lower bound, and compare segmentations that share the nuclear counts; a method that also changes the nucleus, as Proseg does, needs an external reference. The kept ratio measures the error that the mask adds to a nuclear label, not the total error: at equal depth, half B itself recalled only about a third to three quarters of the T cells defined by protein markers (Table 1). The error is not confined to older cell segmentations: stain masks in kidney and synovium kept about three quarters of T/NK labels, and the cells of MERSCOPE and CosMx about half (Table 1). The two renal cell carcinomas, with masks from a segmentation stain and the newest Xenium assay, lost the fewest, but partly because their nuclei carried more of their neighbors’ RNA, which makes the reference less clean rather than the masks better.

Which cells the error reaches determines which conclusions it threatens. Whole-cell counts lost a quarter to half of T/NK labels, but in renal cell carcinoma they recalled more myeloid cells than the nucleus did. Composition dominated by large, RNA-rich classes, therefore, changes little, whereas small or thin cells, such as T/NK, B, mast, and mural cells (Supplementary Table S20), are undercounted, most of all where they sit among larger cells. Claims of immune exclusion, of the depletion of a rare subset, or of a change in its infiltration between conditions are the ones to check against nuclear labels. For large cells, counting noise is the more likely source of label error. A companion study finds that this error, where it confuses subtypes of one lineage, inflates estimates of how a cell’s expression depends on its neighborhood, and that the label and the modeled expression should, therefore, come from different transcripts [30]. The same study finds a third error, systematic error of a labeler that learned a subtype boundary from another labeler’s calls, which reversed niche claims of Niche-DE on semi-synthetic data [30]. Counting noise and segmentation meet in lung cancer labeled from whole cells and from nuclei: nucleus labels in a cell’s own role changed the estimated neighborhood effect as counting noise would, whereas nucleus labels in its neighbors’ role lowered it by more than thinning predicted [30]; Supplementary Table S21 sets the three errors side by side.

We see three practical consequences. First, random relabeling does not bound the real error, because it moves contact statistics in the opposite direction; stress tests should place errors where cells are in contact. The companion study finds the same for niche claims: errors spread at random over subtypes reversed no claim of Niche-DE, whereas a labeler’s own errors did [30]. Contacts are best reported with statistics normalized to the immune class, except where small cells sit inside a layer of larger ones, as T cells in gut epithelium. The direction of the change also depended on the labeling: with the authors’ labels, T/NK contacts with loop of Henle cells in kidney rose rather than fell, and in synovium they barely moved (Supplementary Methods). The direction has to be measured for each labeling, and the test does so. Second, corrections that keep the label cannot, by design, repair it, and models of the neighbors’ RNA trade one error for another. For composition and contacts, we recommend labeling small classes from the nucleus, with the whole cell for cells whose nucleus holds too few counts, and large classes from the whole cell. Third, a contrast between two groups changes with the labeling where the groups differ in exposure. At equal depth, and for classes that mostly lose labels, whole-cell labels scale each group’s count by its kept ratio, so the contrast is biased by the ratio of the groups’ kept ratios, which the labels themselves measure; at full depth, the labelings also differ in which cells they label and how they call regions, so the bias there has to be measured (Supplementary Methods). For contrasts built on counts of lymphocytes within the epithelial layer or of tumor-infiltrating lymphocytes, we recommend nuclear labels, or checking that the groups’ kept ratios agree. That no enrichment call in our data changed sign is what this predicts for claims far from a decision threshold: segmentation error scales magnitudes and contrasts, and it reverses a call only near a threshold or between groups that differ in exposure. Errors that move labels between subtypes of one lineage can do more: in the companion study, a labeler’s systematic errors reversed robust niche claims of Niche-DE on semi-synthetic data [30].

Earlier work measured segmentation admixture in the expression, with the labels held fixed [3, 4, 31]; here we held the counts fixed and followed the labels, cell by cell, without ground-truth labels. Newer methods segment with expression as well as images [32, 33] or recover expression that segmentation leaves unassigned [34], and the kept ratio can measure what they gain. A recent perspective expected rare populations to be the most sensitive to segmentation errors [35]; small cells that border larger ones are such a case, and in ovarian cancer, the confidence of consensus labels fell where the surrounding cell types were most mixed [36]. A generative model of neighbor-derived counts [5] recovered T cells that the nucleus missed but moved errors into other classes, and only protein markers on the same cells revealed both. Assays of RNA and protein on the same cells [37, 38] would allow the same validation on further platforms, as would CosMx data with protein markers that identify individual T cells, which we could not find in public data.

Our study has several limitations. The test sees only the part of the label error that differs between nucleus and whole cell; errors that both share, from the reference, the label granularity, or the classifier, are invisible to it. It covers only cells whose nuclear half holds enough counts to be labeled, about half of the T cells defined by protein markers in renal cell carcinoma. Those it leaves out hold the fewest nuclear counts and are likely among the most exposed, so the gap more likely hides loss than inflates it. The losses are measured at equal depth, which isolates what the mask adds; at full depth, whole-cell counts also label more cells, and in renal cell carcinoma, though not in lung, whole-cell labels then recalled slightly more T cells than nuclear labels (Supplementary Table S16). The nucleus itself collects some of its neighbors’ RNA, about a quarter of the rate of a T/NK cell’s own RNA in the Xenium tissues and more than half in renal cell carcinoma, so the measured loss is a lower bound. Isolated nuclei, which held none of the excess, came from other tumors and were profiled over the whole transcriptome, so that comparison gives a direction rather than a size (Methods). Nuclear and cytoplasmic transcriptomes also differ for biological reasons [39–41], but such differences do not depend on which cell touches which, whereas the excess changes we count went to the contacting neighbor’s type.

Protein markers cover five sections from as many donors on two platforms, the only public imaging transcriptomics data with T-cell protein markers on the imaged cells that we could find (Supplementary Note 2). In the two renal cell carcinomas, more cells were CD3E-positive by protein than are likely to be T cells, but lineage-consistent T cells defined by protein markers gave the same recall ratio (Supplementary Methods), and the partner types in their contact results are approximate, because the kidney reference has no tumor type. Of the lung immunohistochemistry stains, only CD8 and pan-cytokeratin worked reliably, and no number here depends on pan-cytokeratin. The regions in which we placed T cells were defined from their neighbors’ labels; in renal cell carcinoma, regions defined by protein gave the same direction. Finally, each tissue was labeled against one reference, the polygons are two-dimensional, and all analyses are exploratory.

## Methods

### Data

Lung cancer, Xenium: 12 sections of donors L1–L4 on the Lung (289 genes), Custom IO (340), and 5K panels (masks from nucleus expansion for the Lung and Custom IO panels, whose files predate Xenium Onboard Analysis 2.0 and its segmentation stain; from the stain for 5K), with post-Xenium immunohistochemistry (CD8, pan-cytokeratin) of two sections; reference, single-cell Flex of the same donors (35,954 cells; GEO GSE311609) [4]. Gut mucosa (inflammatory bowel disease), Xenium, 32 sections of 15 patients (GSE312420) [16], on four panels, with masks from nucleus expansion or from the segmentation stain (Supplementary Methods). Each patient is one unit, pooled over its sections; 27 sections of 14 patients had at least 1000 cells in the null and entered the extra-change test. Reference, CZ CELLxGENE Census gut. Kidney, Xenium, 20 patients (GSE336890) [17]; reference, Census kidney. Synovium, Xenium Prime 5K, 17 patients with rheumatoid arthritis (Zenodo 15058262) [18]; reference built from GSE200815 [42] (Supplementary Methods). Lung cancer, CosMx, 8 sections of 5 patients, 960-gene panel [19], 171 over-detected probes excluded (Supplementary Note 1). Renal cell carcinoma, Xenium Protein (10x Genomics, CC BY 4.0), one FFPE section of each of two donors with 27 proteins on the same cells and the Human Multi-Tissue and Cancer panel (377 genes): donor 1, renal cell carcinoma [20]; donor 2, clear cell renal cell carcinoma [21]. Reference for both, Census kidney, which has no tumor type. Uterine cancer, MERSCOPE [22]: one section, 500-gene panel, 758,506 cells segmented by the platform, and an antibody against CD45RO imaged in the same run; reference, Census uterus (normal donors; no tumor cells). Liver, CosMx, metabolic dysfunction-associated steatotic liver disease (GSE312698) [43]; reference, Census liver. Panels, software versions, and the preparation of each dataset are in Supplementary Methods. Units of inference are donors or patients (liver: tissue pieces); downloads were not modified.

### Labels and count sets

Multinomial maximum likelihood of a cell’s counts against the mean gene proportions of each reference type, with a 1% flat floor, a uniform prior (sensitivity: floor 0.1, 1, and 5%, and a class prior fitted per section and count set by expectation maximization) and, in lung and CosMx, a pergene platform factor refitted per section; cells with fewer than ten counts were unlabeled. Second labeler: CellTypist [44] trained on the same reference cells, thinned to 20–200 counts. Count sets per cell: nuclear counts; halves A and B (binomial, *p* = 0.5); whole-cell counts minus A thinned to the depth of B (wholeB), at other depths, or not thinned; counts outside the nucleus thinned to the depth of B (cytoB).

#### Label audit

For each section and for the nuclear and whole-cell labels, we listed genes detected far above their share in every reference type and counted the calls that exist only through such a gene (claimbreak compartment audit; Supplementary Methods). Synovium was relabeled without CCL5 and EEF1G, the two genes the audit found deciding calls in the first labeling.

### Extra-change test

Contact neighbors: up to six nearest cells within 30 µm, typed by nuclear labels. Shares of a type among neighboring cells count only labeled cells. Contact neighbors are the nearest cells, labeled or not; an unlabeled neighbor has no type. Contact readouts and colocalization use only the cells labeled by both labelings compared. Expected share *e*: for each label change *a → b*, the share of all eligible cells labeled *a* that touch a type-*b* cell, averaged over the changes (the share if a cell’s chance of changing to *b* did not depend on contacting *b*). Extra-change ratio: a label change is a hit when its new type is that of a contact neighbor; with *H* the number of hits and *E* the number expected if changes ignored the neighbors (the sum of *e* over the changes), each per cell compared, in the test (*T*) and the null (0), the ratio is (*H_T_ −H*_0_)*/*(*E_T_ −E*_0_): observed over expected hits among the extra changes, pooled per unit over sections (sums of numerators and denominators). Units are evaluable with at least 50 extra changes and a denominator positive in more than 90% of 2000 parametric bootstrap draws. Block bootstrap: the same sums per 200-µm tile, tiles of a unit resampled with replacement (1000 draws). Margin sensitivity: cells whose half A label beats the second-best type by less than 5, 10, or 20 nats removed (at 20, 0.38–0.51 of cells kept). Pair excess: rate of *a → b* at contact minus absent, minus the same contrast in the null. Confirmed labels: nuclear counts split into three multinomial thirds A, B, C; test, whole-cell counts minus A and C thinned to the depth of B; null, B against A; only cells whose label from A equals that from C (claimbreak compartment compare --agree). Boundary contact: cells whose deposited segmentation boundaries come within 1 µm; far-only neighbors then exclude contacting cells. Matched expected share (claimbreak 1.7.81): for each change *a → b*, the share of the other eligible type-*a* cells that touch a type-*b* cell among those with the same regional share of *b* (cells whose centers lie 30–100 µm away, counted on a 5-µm grid; 10 bins: none, eight quantiles of the positive shares, no cell in the ring) and the same number of contacts; strata with fewer than 20 other cells fall back to the regional bin alone, then to the section. The matched ratio is conservative: where contacting and regional composition agree, it credits the region. Its check on simulated sections is in Supplementary Methods.

### Masks and exposure

Segmentation series: each transcript outside a nucleus given to the cell whose nucleus boundary is nearest, within *d* = 1–15 µm; nuclear counts identical in every mask. Cell dimensions (sizes.py, per section and nuclear label class): nucleus and deposited-mask areas as equivalent-circle diameters; gap, the centroid distance minus both equivalent nuclear radii, to the nearest and the sixth-nearest cell (half the gap to the nearest nucleus was 0.3–1.4 µm, all labeled cells); growth, the share of the 15-µm mask’s counts outside the nucleus held at each expansion. Border share from polygons (lung, gut, kidney, renal cell carcinoma): each deposited cell polygon densified to points every 1 µm; a point is held by the nearest point of another cell’s polygon within 1 µm, and a neighbor’s share is the fraction of the cell’s points it holds (points held by no cell count as free outline, so the shares of a cell’s neighbors sum to its contacted fraction; claimbreak compartment exposure --boundaries). Nuclear-disk proxy (synovium, CosMx, and the neighbor-aware analyses): nuclei as disks of their measured area; gap = center distance minus both radii; masks touch for gaps up to 10 µm; the cell’s circumference in 72 angular bins, each given to the contacting neighbor whose disk covers it with the smallest gap. Kept ratio: cells whose half A label is T/NK and that keep it in the test, over those that keep it in the null, per stratum of border share, outside share, or regional composition (share of a type among cells within 20 µm), pooled per unit. The same ratio for every other class was computed from the stored labels (small_classes.py, units with at least 100 cells kept in half B); for T/NK cells it gave 0.71 in gut, 0.76 in kidney, 0.76 in synovium, and 0.57 in lung, close to the main analysis (Table 1). Destinations: excess rate *a → b* over the null against *b*’s border share (claimbreak compartment exposure).

### Foreign load

Marker genes: highest mean reference proportion in one type, at least four times the next. Type from one nuclear half; foreign share: counts in other types’ markers over all marker counts, pooled over cells. *ρ*: foreign over own marker counts. For Supplementary Fig. S2, whole-cell counts were split binomially; *ρ* from one half, the label from the other thinned to the depth of B.

### Neighbor RNA by contact

Cell types came from the nuclear half A labels. The partner was the tumor in lung, epithelial cells in gut, fibroblasts in synovium, and proximal tubule in kidney. Per section, partner markers were chosen from nuclear counts without positions: the 50 genes most expressed in partner cells among genes detected in at least 20% of them and at least ten times higher, as mean counts, in partner than in T/NK cells; T/NK markers the same way. The load is marker counts per 1000 counts of a cell, from its whole-cell and from its nuclear counts. Contact is a partner cell among the six nearest cells within 30 µm. Per section with at least five cells at contact and five away, the load at contact was divided by the load away; ratios are medians over sections, with 95% intervals from a bootstrap over donors or patients. The share of the excess held by the nucleus is the median over sections of the contact-minus-away load in nuclear counts over that in whole-cell counts. The reverse direction exchanges the roles. Slide-GoTags (Broad Single Cell Portal SCP3667 [23]): raw counts of the deposited matrix, the deposited cell types, positions in micrometers, and the same rules, with tumor markers also detected in at most half of T nuclei. Three things would pull the isolated-nuclei ratio toward one: nuclei that are neighbors in a 20 µm section but never touched, unmapped nuclei that remove true partners, and the removal of nuclei with high ambient RNA before cell typing; the test with cells removed covers only the second. Protein, renal cell carcinoma: T cells CD3E^+^ CD45^+^ CD68*^−^* PanCK*^−^* and, as partners, tumor cells PanCK^+^ (donor 2) or CD45-negative cells (donor 1, where PanCK does not separate) by the protein calls (Protein), markers chosen from nuclear counts for these classes, and 500 µm tiles of the section as units. The positional and mislabeling checks are in Supplementary Methods.

### Transcript-based segmentation

Proseg 3.2.0 [6] was run on the Xenium transcripts of L1 and L3; each Proseg cell was mapped to the Xenium nucleus whose transcripts it holds most of, its nuclear counts are its transcripts over that nucleus, and it was labeled with the same classifier and reference and compared with the deposited masks on the same cells. The same was done for the two renal cell carcinoma sections (counts from transcripts with quality value at least 20), with the options of their deposited-mask labels, and the deposited labels of the mapped cells were scored again on those cells. Baysor 0.9.0 [2] was run with its Xenium preset (at least 50 molecules per cell, prior confidence 0.5, the Xenium cells as prior) on tiles of 1 mm with a 30-µm overlap; each Baysor cell was tied to the Xenium nucleus whose transcripts it held most of and kept only in the tile whose core holds that nucleus (claimbreak baysor_tiles.py), then labeled and compared like the Proseg cells. The numbers of mapped cells are in Supplementary Methods.

### Protein markers

Multiplex IHC (DAPI, three Opal channels, autofluorescence; 0.496 µm per pixel) of the Xenium sections of L1 and L3. DAPI maxima registered to Xenium nuclei (density cross-correlation, mutual nearest neighbors, affine refit; matches within 2 µm). Positivity by mixture model on the 2–5-µm ring intensity after autofluorescence subtraction; CD8 channel assigned by its enrichment in T/NK over tumor labels. Recall: share of CD8^+^ cells labeled T/NK. Chance matches: *f*, the matched share at 2 µm under 40 registrations shifted by 50–300 µm over the matched share of the registration; corrected recall (*r−fp*)*/*(1*−f*), with *p* the share of matched cells labeled T/NK in the same count set (an approximation of the T/NK share among chance matches). Renal cell carcinoma: background-subtracted protein intensity per cell (Xenium Onboard Analysis); positive cells from a two- or three-component mixture on log(1 + *x*) of the cells with non-zero intensity (highest component, Ashman *D ≥* 2); T = CD3E^+^, CD8 = CD8A^+^, CD4 = CD4^+^CD3E^+^, B = CD20^+^, myeloid = CD68^+^; contact with a CD68^+^ cell among the six nearest within 30 µm (pan-cytokeratin did not separate, *D* = 1.91); loss 1*−r*_wholeB_*/r*_nucB_ with 95% intervals from resampled 200-µm tiles (1622 and 2707 tiles; 1000 draws; claimbreak xenium_protein.py). Non-immune share: the fraction of a cell’s polygon outline (as above) held by cells that are not CD45^+^ by protein, binned at 0, 0.25, 0.5, 0.75, and 1; loss per bin with block intervals, and a least-squares slope of the loss on the bin midpoints weighted by the square root of the cells (bins of at least 200 cells; xenium_protein.py border--base outline). MERSCOPE: nuclei segmented from the DAPI image with StarDist [45], each assigned to the cell whose outline in the same plane contains its centroid; a transcript is nuclear when it lies in a nucleus of its own cell, and cells without a nucleus (0.68 of cells own one) leave the nuclear count sets. CD45RO: the high-pass signal per µm^3^ of cell volume, positive cells by the mixture rule above, read as the T class; CD45RO also marks some activated myeloid cells, which lowers recall in every count set alike but not the recall ratio. The checks that the markers identify individual cells, the published labels of donor 2, and the MERSCOPE segmentation settings are in Supplementary Methods.

### Labels against protein markers

For each section with protein markers and each segmentation, two ratios were generated relative to nuclear half B at equal depth. Label only: the kept ratio of a class (cells whose half A label is the class and that keep it in the test, over those that keep it in half B; claimbreak compartment summary, units_kept.csv). Protein marker: the recall of protein marker-positive cells of the class by the test counts over their recall by half B (lung, CD8^+^ cells by immunohistochemistry, any CD8 call; renal cell carcinoma, CD3E^+^ T, CD20^+^ B, and CD68^+^ myeloid cells). Comparisons: the expansion series (*d* = 1–15 µm, whole-cell counts minus A thinned to B), the deposited masks, and Proseg in L1 and L3; whole-cell and outside-nucleus counts in the two renal cell carcinoma sections, for T, B, and myeloid cells, and with Proseg, for T cells. The 30 comparisons of T and B cells are nine per lung section (the nucleus expansions, the deposited masks, and Proseg, all for CD8^+^ T cells) and six per renal cell carcinoma section (T and B cells from whole-cell and outside-nucleus counts, and T cells from Proseg’s whole-cell and outside-nucleus counts); myeloid cells, which gain labels, are shown but not counted, and the Baysor comparisons are reported separately. Agreement over the T and B comparisons: mean and largest absolute difference, per donor and over all comparisons. Agreement is summarized per donor because the comparisons of one donor share cells. Why the two ratios agree, and when they do not, is derived in Supplementary Methods, with a ground-truth simulation of both cases. Recall counts the protein-positive cells labeled in a count set; where stated, unlabeled cells count as missed (Supplementary Methods). Hybrid labels (xenium_protein.py and ihc_truth.py truth --hybrid nucleus:whole): the nuclear label where the nucleus held at least ten counts, else the whole-cell label, scored in the same way. Strict protein classes, the cell-specificity of the protein classes, and the synovium outcomes are in Supplementary Methods.

### Label loss by region

For each cell labeled as the immune type by nuclear half A: region share, the share of partner cells among the other cells with a half A label within 25 µm; bins away (< 0.2), border (0.2–0.5), and inside (*≥* 0.5). Partners: epithelium (gut), tumor (lung), fibroblasts (synovium), and in kidney the loop of Henle, the proximal tubule, or every non-immune label. Loss, one minus the kept ratio of whole-cell counts at the depth of half B over half B, per bin, pooled over the sections of each unit, bins of at least 20 cells; medians over units, by mask type, and per unit inside minus away (claimbreak region_loss.py, region_loss.zsh). Protein truth, renal cell carcinomas: T cells by CD3E protein (and the strict T class, Supplementary Methods), region cells CD45-negative or PanCK-positive by protein, the same region share and bins; loss of whole-cell labels at the depth of half B relative to half B, with paired 95% tile-bootstrap intervals (200 µm tiles, 1000 draws; protein_region.py). The identity and state of the calls, and the counts of T cells among epithelial cells by condition, are in Supplementary Methods and Supplementary Table S18.

### Contact statistics and random relabeling

Contacts per 100 partner cells: immune cells contacting the partner per 100 partner cells; enrichment calls: permutation *z*, *|z| ≥* 2 and | log_2_ *O/E| ≥* 0.2. Random relabeling: 5, 10, or 15% of cells reassigned (ten draws), the new type drawn uniformly or from the section’s composition with-out the cell’s own type (composition preserved in expectation); for contacts per 100 partner cells applied to the whole-cell labels and compared with them, and to the nuclear labels where stated. Same changes placed at random: each half B label replaced by a draw from the section’s confusion row P(whole-cell label | half B label), ten draws, compared with the half B labels (claimbreak compartment contacts --confusion; per100_conf.zsh); with the partners fixed, every partner cell kept at its half B label in the real and in the random labels, so that only the immune side changes (--fix-partners; per100_fixed.zsh); renal cell carcinomas, with partners typed by protein, fixed at their non-immune half B labels, or typed by each labeling (protein_contacts.py--confusion; protein_position.zsh). Placement of the real changes and contacts scored against protein: Supplementary Methods.

### Corrections

RCTD (spacexr 2.2.1, doublet mode), SPLIT 0.3.0 (default, spatially aware, SPLIT-shift), and cellAdmix 0.1.0 on 30,000-cell windows of the five Lung-panel sections; corrected counts relabeled with the same classifier. Absorbed cells: immune by nuclear label and labeled a contacting nonimmune type from whole-cell counts.

### Neighbor-aware labeling

For candidate type *k*, log *L*(*k*) = ∑*_g_ n_g_* log *q_kg_* +∑*_g_ o_g_* log[(1*−ψ*)*q_kg_* +*ψc_g_*], with *n* and *o* the nuclear and outside counts, *q_k_* the type profile, and *c* the expected contamination: the reference profiles of the neighbors’ nuclear types weighted by their RNA and by their border share (contacting), their mean RNA within 20 µm, and the section composition (weights 1, 0.5, 0.1, set by hand, not fitted; sensitivity in renal cell carcinoma: contact only, and the regional and ambient weights halved or doubled, claimbreak xp_batch.zsh remsens; results in Supplementary Table S15). The foreign share *ψ* = *κE/*(1 + *κE*) follows the cell’s exposure *E* (neighbor over own RNA), with one *κ* per section fitted by maximum likelihood (claimbreak decontam). Evaluated without shared counts: labels from half B plus the outside counts against labels from half A; *κ* is fitted on the counts of the labeled count set itself (half B plus outside), never on half A. The count set with the whole nucleus contains half A and is evaluated against protein only.

### resolVI

resolVI (scvi-tools 1.5.1) was run on square windows of about 30,000 cells, those per renal cell carcinoma section with the most cells labeled T/NK from whole-cell counts. Following the authors’ relabeling, we trained an unsupervised model on the whole-cell counts, built a nearest-neighbor graph in its latent space, and set to unknown the label of each cell whose neighbors carried other labels in more than 5% of its connectivity (the authors’ threshold for coarse labels). A semi-supervised model trained on the kept labels then relabeled every cell (claimbreak resolvi_rcc.py). The nuclear, whole-cell and resolVI labels of the same cells were scored against protein as above, with tile-bootstrap intervals for paired differences. Choosing windows rich in T/NK cells favors gains in T recall. Window sizes, training settings, and the sensitivity runs are in Supplementary Methods.

### Use of large language models

The authors used Claude (Anthropic; several model versions, 2026) as an assistant throughout the study: to draft and revise analysis code, the claimbreak package and its tests, and manuscript text; to check the text against the computed numbers; and to propose analyses and critique drafts. All analyses were run by the authors on their own computers. No number in this manuscript was produced by the language model: every value is inserted from numbers files computed from the result folders. The authors reviewed, tested, and edited all code and text and take full responsibility for the content of this article.

### Statistics and software

Summaries are medians over units with ranges, and the number of units in which an effect has the stated sign. Analyses are exploratory; no test has a pass or fail threshold. We used claim-break 1.7.103 (Code availability); the results were produced with development versions up to 1.7.101, and later versions add one example script and do not change them. Scripts named in Methods and Supplementary Methods are claimbreak tools (the package’s command-line interface or its examples folder), except sizes.py, small_classes.py, ibd_bias.py, ibd_decomp.py and ibd_histology.py, which are among our analysis scripts (Code availability). Every number in the text is inserted from numbers files computed from the result folders; the result tables and the script that fills the numbers into the text will be deposited on publication (Data availability).

## Data availability

All data analyzed here are publicly available; one dataset requires registration (below). Lung cancer, Xenium, single-cell Flex, and post-Xenium immunohistochemistry: GEO GSE311609 [4]. Gut mucosa: GSE312420 [16]. Kidney: GSE336890 [17]. Synovium: Zenodo 15058262 [18]; synovium reference, GSE200815 [42]. Lung cancer, CosMx: the public CosMx non-small-cell lung cancer dataset [19]. Liver, CosMx: GSE312698 [43]. Breast cancer, CosMx same-slide RNA and protein (assessed for protein truth only): Bruker Spatial Biology [46]. Head and neck carcinoma, CosMx (stains only): Dryad doi:10.5061/dryad.n2z34tn7x [47]. Tumors profiled with Slide-GoTags (iso-lated nuclei): Broad Single Cell Portal SCP3667 [23]. Renal cell carcinoma, Xenium Protein: 10x Genomics datasets [20, 21] (CC BY 4.0). Uterine cancer, MERSCOPE with CD45RO co-staining: Vizgen Data Release Program [22], available from Vizgen after registration; we do not redistribute these data. References: CZ CELLxGENE Census. The result tables behind every number and figure, the numbers files, and the script that fills every number into the text are available from the corresponding author on request and will be deposited at Zenodo on publication; data from the Vizgen Data Release Program will not be included.

## Code availability

claimbreak (version 1.7.103, https://github.com/velgart/claimbreak; archived at doi:10.5281/zenodo.23987641; BSD 3-Clause license) runs every analysis; claimbreak kept-ratio runs the label-only check, with a single-cell reference, on a user’s own Xenium or CosMx data; for MER-SCOPE, the example script merscope_nuclei.py first segments nuclei from the DAPI image. The two packages of this work have different jobs: claimbreak measures how segmentation changes the labels, and nichepeek [48] measures how label error biases estimates of neighborhood effects [30]. The analysis scripts that computed the result tables and figures are available to editors and reviewers at submission and from the authors on request.

## Acknowledgements

This work was supported in part by NIH grants R01 HL155107 and R01 HL166137; by AHA MERIT grant 1185447; by EU Horizon Health 2021 grant 101057619; and by a Ruggeri Family Network Medicine grant to J.L. The authors wish to thank Rose Cannistraro for expert technical assistance.

## Author contributions

VE conceived the study, developed the method and the claimbreak software, performed the analyses, and wrote the manuscript. JL conceived and supervised the study, contributed to the interpretation of the results, and wrote the manuscript. Both authors read and approved the final manuscript.

## Supplementary Information

**Table S1.** How label-based spatial methods treat label error. Methods whose robustness test reassigns labels independently of position, and corrections that keep the labels; examples, not a systematic review.

| work | domain | label error tested |
|---|---|---|
| RobustCCC [7] | single-cell CCC | labels permuted uniformly at random in 5–15% of cells |
| HRCHY-CytoCommunity [9] | spatial niches | 10–50% of labels perturbed at random; structures stable up to 40% |
| scNiche [8] | spatial niches | a random share of labels set to “ambiguous” |
| QUICHE [49] | spatial niches, protein imaging | none on labels; settings varied on simulations with correct labels; null shuffles types at random |
| cellAdmix [3] | imaging transcriptomics | none; admixed molecules removed with labels fixed |
| SPLIT [4] | Xenium | none as a test; one CD8<sup>+</sup> cell annotated as tumor; SPLIT-shift swaps primary and secondary types |
| resolVI [5] | imaging transcriptomics | none as a test; cells relabeled by a classifier trained on the existing labels; cells whose label differs from the original annotation shown |
| STARLING [50] | imaging mass cytometry | segmentation errors favor neighbors’ clusters |
| Xenium best practices [13] | Xenium segmentation and preprocessing | none on labels; cells of each new segmentation typed by majority vote from the nuclear annotation; expression purity scored; mis-segmentation simulated without positions |
| segmentation benchmark [14] | five platforms | none on labels; neighbor markers in expression scored with labels as given |
| annotation and power benchmarks [51–53] | spatial | none; labels correct by construction |

### Supplementary Note 1: CosMx probe exclusion

With all panel genes, labels of the three serial sections of patient Lung5 differed by up to 0.24 in the share of one class and one section labeled most cells as vessel. 146–207 probes per section were detected at more than twice their largest share in any reference type, mostly the same in every section. Excluding the 171 probes listed in at least half of the sections (a rule that uses no labels and no compartment information), the Lung5 sections differed by at most 0.08 and their nuclear halves agreed on 0.77–0.81 of cells.

### Supplementary Note 2: liver, a dataset whose labels fail first

#### Summary

In liver biopsies and explants with metabolic dysfunction-associated steatotic liver disease, on CosMx with a 1000-gene panel [43] (499,417 cells in 11 tissue pieces), the classifier settings used for lung collapsed the labels of 7 pieces onto a single class that is a minority in liver: 84–94% of nuclear labels were cholangiocyte and 66–93% of whole-cell labels endothelial. The nuclear halves of these pieces still agreed in up to 0.94 of cells because both halves were given the same incorrect class. Without the per-gene platform factor, no class exceeded 81% of any piece, and the agreement of the halves separated the slides: 0.85–0.91 in the two explant sections, 0.67–0.75 in the biopsies of one slide, and 0.30–0.49 in those of the other two. On the 4 pieces whose halves agreed in at least 0.6 of cells (the lowest agreement of the Xenium sections was 0.56, over 83 sections; one section below 0.6, in kidney), whole-cell labels kept 0.40–0.70 of the T/NK labels of the null at equal depth (intervals below one in 3 of 4; in the largest explant section 0.59, 0.55–0.64) and lowered T/NK, B, and myeloid contacts per 100 hepatocytes in 12 of 12 piece–type pairs. Liver, therefore, supports the loss of T/NK labels but adds little to the contact test.

**Table S2.** The extra-change test under alternative choices. Extra-change ratio at contact (whole-cell counts at the depth of half B against half A, over the null). Main analysis: pooled per unit, range over evaluable units (lung, gut, kidney, synovium, CosMx) or per section with its 95% block interval (renal cell carcinoma). Confident cells: cells whose half A label beats the second-best type by less than 20 nats removed. Confirmed labels: only cells whose labels from two independent thirds of the nucleus agree. Boundary contact: neighbors that share a segmentation boundary instead of the six nearest cells. Proseg and Baysor: the same sections resegmented, on the cells each keeps (lung L1 and L3; renal cell carcinoma 1 and 2). Values for the variants are pooled ratios, medians over units. Matched on the composition of cells 30–100 µm away and on the number of contacts (Methods), the ratio had medians of 1.25–1.54 in the four Xenium tissues, with block intervals above one in 45 of 51 donors and patients (47 unmatched); matching kept 0.59–0.76 of the excess over one. With confirmed labels, block intervals were above one in 50 of 52 evaluable units; block intervals were 0.8–1.2 times as wide as parametric ones and excluded one in the same units. A second labeler, CellTypist [44]: 42 of 55 units evaluable, all above one. Classifier settings (flat floor 0.1, 1, or 5%; uniform or fitted class prior): the T/NK kept ratio was 0.51–0.59 in lung and 0.70–0.73 in gut (medians over units, range over the six settings), and T/NK per 100 partner cells fell by 44–53% and 46–49% in every unit.

|  | main analysis | confident cells | confirmed labels<br>(all → confirmed) | boundary contact<br>(nearest → boundary) | Proseg | Baysor |
|---|---|---|---|---|---|---|
| lung | 1.48–2.14 | 1.74 | 1.79 → 1.77 | 1.76 → 1.81 | 1.38, 1.46 | 1.41, 1.44 |
| gut | 1.47–2.44 | 1.74 | 1.74 → 1.61 | 1.83 → 1.92 | — | — |
| kidney | 1.08–1.98 | 1.48 | 1.56 → 1.43 | 1.64 → 2.09 | — | — |
| synovium | 1.05–1.87 | 1.30 | 1.43 → 1.33 | — | — | — |
| lung, CosMx | 1.41–2.35 | 1.61 | 1.69 → 1.62 | — | — | — |
| renal cell carcinoma 1 | 1.34 (1.30–1.37) | — | — | 1.34 → 1.55 | 1.26 | 1.22 |
| renal cell carcinoma 2 | 3.20 (2.95–3.50) | — | — | 3.20 → 5.12 | 2.36 | 2.33 |

**Figure S1.**
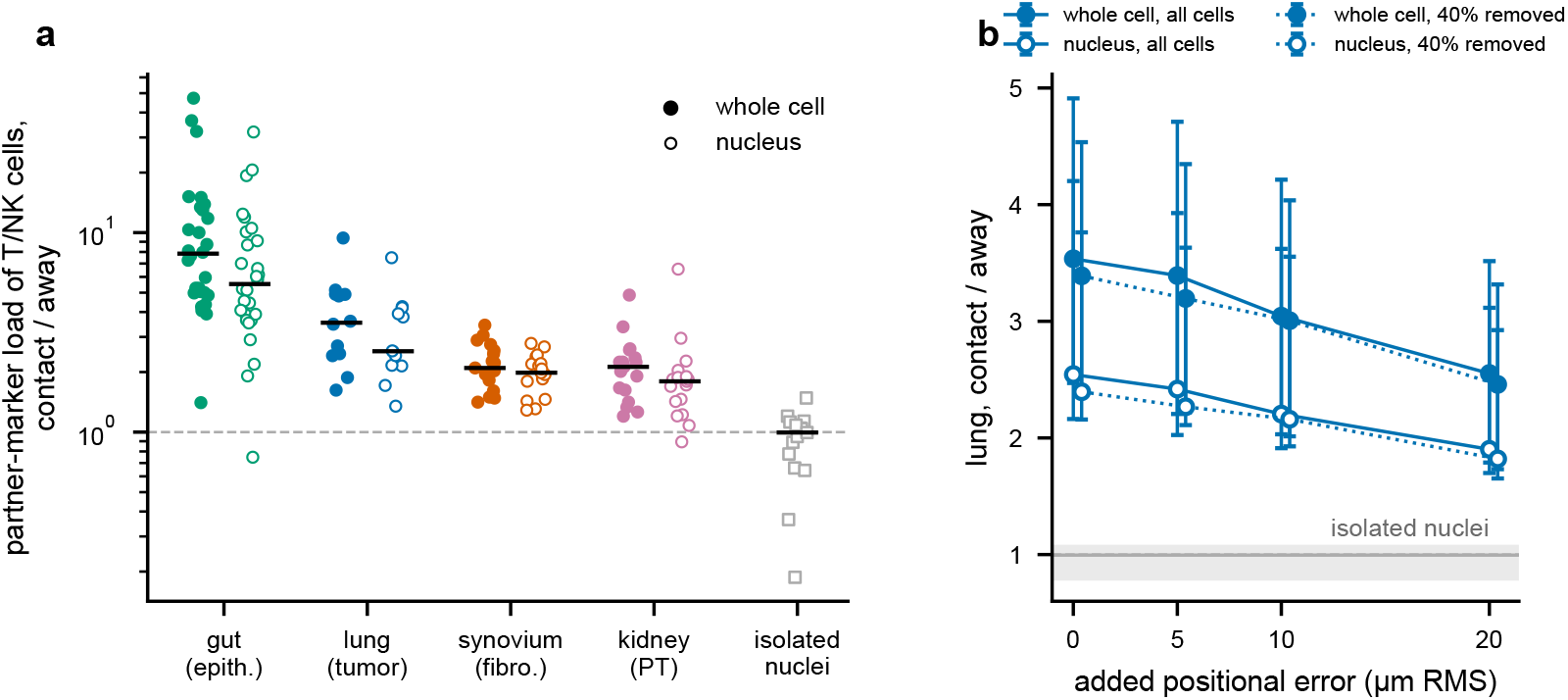
The neighbor’s RNA by contact in whole cells, imaged nuclei, and isolated nuclei. a, Per section or crop, the partner-marker load of T/NK cells at contact (a partner among the six nearest cells within 30 µm) over that of T/NK cells away from one, from whole-cell (filled) and nuclear counts (open) of the same cells, four Xenium tissues; squares, T nuclei next to tumor nuclei over T nuclei away from them in tumors profiled with Slide-GoTags (isolated nuclei); bars, medians; dashed, one. b, Lung: the same ratio after moving every cell by a random positional error (root mean square), with all cells (solid) and with 40% of cells removed (dotted); medians over sections with 95% bootstrap intervals over donors; gray, the Slide-GoTags median and interval.

**Table S3.** Synovium: expected and observed. Expected ranges written from lung, gut, and kidney before the synovium sections were labeled. Observed, medians over patients (range), Harmony-corrected reference, for the first labeling, which an over-detected probe (CCL5) misled (Supplementary Methods, label audit), and for the audited labeling without CCL5 and EEF1G, used everywhere else in the paper. E, outcomes that follow from the measurement; T, outcomes that depend on tissue architecture. E3*^′^*: patients with at least 100 T/NK cells kept in nuclear half B (16 of 17). E4: patients with a ratio of at least one in brackets. T3: split typing, as every foreign-share value, and without it, the rule of our first evaluation. Foreign shares and T3 without the genes each labeling excluded.

|  | expected | first labeling | audited labeling |
|---|---|---|---|
| E1 T/NK, half B / half A (noise) | 0.97–1.03 | 1.00 | 1.00 |
| E2 extra-change ratio at contact | > 1; about 1.3–1.7 | 1.47, interval > 1 in 14 of 15 | 1.45 (1.05–1.87), interval > 1 in 15 of 17 |
| E3 T/NK, whole cell / half B (label composition) | about 0.65–0.80 | 0.65 | 0.75 (0.65–0.85), 17/17 below one |
| E3′ T/NK kept ratio, whole cell / half B | — | 0.72 | 0.77 (0.66–0.87), 16/16 below one |
| E4 large types hold or gain, whole cell / half B | ≥ 1; T/NK loses most, B less | fibroblast 1.12 (17), myeloid 1.07 (13), mural 1.09 (16), endothelial 1.05 (14), plasma 1.19 (15), B 0.95 (4); T/NK 0.65; of 17 | fibroblast 1.10 (17), myeloid 0.99 (8), mural 0.94 (2), endothelial 0.98 (3), plasma 1.03 (10), B 0.82 (0); T/NK 0.75; of 17 |
| E5 T/NK foreign share, nucleus → cytoplasm | about +0.3 | 0.30 → 0.67 | 0.40 → 0.71 |
| E6 random 10% raises T/NK per 100 fibroblasts | every patient | 17/17 (+20%) | 17/17 (+28%) |
| T1 lost T/NK at fibroblast contact go to | fibroblast, myeloid | fibroblast 0.42, myeloid 0.26 | fibroblast 0.25, myeloid 0.37 |
| T2 extra loss at fibroblast contact | 0 to +0.05 | +0.073 | +0.066 (17 of 17 above zero) |
| T3 T/NK cells with more foreign than own marker counts | 0.2–0.4 | 0.30 (0.19–0.48); without split typing 0.26 | 0.55 (0.35–0.68); without split typing 0.55 |

**Table S4.** The overlap control. Ratios of cells given the class by whole-cell counts over those given it by nuclear half B (equal depth), on cells labeled in A, B, and the whole cell, with 95% block intervals over 200-µm tiles. Kept, cells labeled with the class from half A, split by protein; recall, protein-positive cells, split by the half A label. Renal cell carcinoma, protein T (CD3E^+^); lung, CD8^+^ by immunohistochemistry (any call), against T/NK labels. Overlap: share of protein-positive cells labeled with the class from half A. B cells of renal cell carcinoma donor 1 were the one exception to the third column: whole-cell counts gave the B label to 1.11 (1.02–1.20) times as many protein B cells that half A had labeled otherwise as half B did (1422 cells; donor 2, 0.93).

|  | kept, protein<sup>+</sup> | kept, protein<sup>−</sup> | recall, half A other type | overlap |
|---|---|---|---|---|
| renal cell carcinoma 1 | 0.92 (0.91–0.92) | 0.78 (0.76–0.80) | 0.87 (0.85–0.88) | 0.63 |
| renal cell carcinoma 2 | 0.96 (0.95–0.96) | 0.90 (0.88–0.92) | 0.94 (0.93–0.96) | 0.62 |
| lung L1 | 0.70 (0.66–0.74) | 0.60 (0.56–0.65) | 0.54 (0.38–0.72) | 0.45 |
| lung L3 | 0.54 (0.52–0.56) | 0.47 (0.45–0.48) | 0.28 (0.23–0.33) | 0.72 |

**Table S5.** Protein marker CD8^+^ cells labeled T/NK (lung; matches within 2 µm). Protein marker CD8^+^ cells matched and labeled: 1471 (L1) and 11,065 (L3). A share *f* of matches is expected by chance (0.32 in L1, 0.19 in L3); corrected recall (*r − fp*)*/*(1 *− f*) with *p* the T/NK share of matched cells in each count set (Methods). Corrections were run on 30,000-cell windows and are compared with the uncorrected labels of the same windows. Proseg rows: cells mapped to Xenium nuclei (Methods).

| count set | recall |  | chance-corrected |  |
|---|---|---|---|---|
|  | L1 | L3 | L1 | L3 |
|---|---|---|---|---|
| nucleus | 0.46 | 0.72 | 0.60 | 0.81 |
| nuclear half B | 0.46 | 0.72 | 0.62 | 0.81 |
| whole cell (deposited mask) | 0.32 | 0.42 | 0.42 | 0.48 |
| whole cell, equal depth | 0.31 | 0.37 | 0.41 | 0.42 |
| windows, uncorrected | 0.33 | 0.37 | 0.44 | 0.42 |
| SPLIT | 0.27 | 0.23 | 0.36 | 0.26 |
| RCTD | 0.27 | 0.22 | 0.37 | 0.26 |
| SPLIT-shift | 0.33 | 0.23 | 0.45 | 0.26 |
| cellAdmix | 0.33 | 0.38 | 0.44 | 0.43 |
| half B plus outside counts | 0.26 | 0.30 | 0.35 | 0.34 |
| the same, neighbor-aware | 0.32 | 0.47 | 0.42 | 0.53 |
| whole cell, neighbor-aware (contains the nucleus) | 0.38 | 0.58 | 0.50 | 0.65 |
| Proseg, nuclear half B | 0.48 | 0.75 | 0.65 | 0.84 |
| Proseg, whole cell | 0.41 | 0.65 | 0.54 | 0.73 |
| Proseg, whole cell, equal depth | 0.41 | 0.64 | 0.56 | 0.72 |

**Table S6.** Labels given to protein marker CD8^+^ cells (lung L3). Share of matched CD8^+^ cells given each label by nuclear and by whole-cell (deposited mask) counts; the lost T/NK labels went to the types around the cells.

| label | nucleus | whole cell |
|---|---|---|
| fibroblast and muscle | 0.12 | 0.29 |
| myeloid | 0.10 | 0.17 |
| tumor | 0.006 | 0.05 |

**Figure S2.**
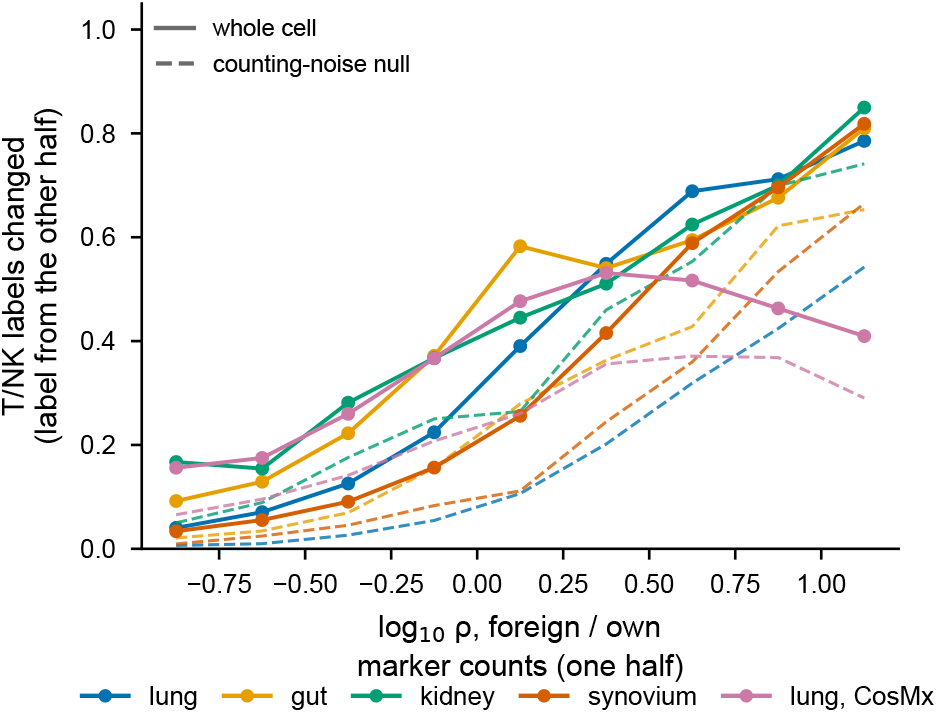
Foreign load predicts the label change. Share of T/NK labels that change (solid) and the counting-noise null (dashed) against *ρ*, foreign over own marker counts, with *ρ* from one binomial half of the whole-cell counts and the label from the other.

**Table S7.** Renal cell carcinoma: protein-defined cells given the matching label. Two donors, one section each. Recall from nuclear half B and from whole-cell counts at the same depth, and the loss, overall and by contact with a CD68^+^ cell (typed by protein). Cells: protein-positive cells of the class with a half B label, the denominator of the recalls. Positive shares: over all cells in the label table that have protein values, labeled or not. Donor 1: 465,534 such cells; positive shares CD3E 0.33, CD8A 0.19, CD68 0.20, CD20 0.055. Donor 2: 690,322 such cells; CD3E 0.33, CD8A 0.15, CD68 0.12, CD20 0.041; pan-cytokeratin separated (*D* = 2.26), but contact is reported for CD68 in both. Nuclear labels recalled 0.61 of T cells at CD68^+^ contact and 0.65 without in donor 1, 0.64 and 0.58 in donor 2. At PanCK^+^ contact in donor 2, nuclear labels recalled 0.58 of T cells, against 0.61 without, and whole-cell labels at the same depth 0.54 against 0.59.

| protein class | cells | half B | whole cell | loss (%) | at CD68<sup>+</sup> contact | without |
|---|---|---|---|---|---|---|
| *donor 1, renal cell carcinoma* |  |  |  |  |  |  |
| T (CD3E<sup>+</sup>) | 83,529 | 0.63 | 0.57 | 9.2 | 11.4 | 6.2 |
| CD8 (CD8A<sup>+</sup>) | 48,778 | 0.74 | 0.66 | 9.9 | 11.5 | 6.8 |
| CD4 (CD4<sup>+</sup>CD3E<sup>+</sup>) | 16,153 | 0.77 | 0.71 | 7.4 | 11.0 | 5.2 |
| B (CD20<sup>+</sup>) | 15,187 | 0.87 | 0.86 | 1.6 | 6.2 | 1.1 |
| myeloid (CD68<sup>+</sup>) | 54,899 | 0.82 | 0.85 | −3.7 | −4.0 | −2.0 |
| *donor 2, clear cell renal cell carcinoma* |  |  |  |  |  |  |
| T (CD3E<sup>+</sup>) | 110,950 | 0.61 | 0.59 | 3.9 | 5.0 | 2.7 |
| CD8 (CD8A<sup>+</sup>) | 47,706 | 0.73 | 0.69 | 4.5 | 5.4 | 3.5 |
| CD4 (CD4<sup>+</sup>CD3E<sup>+</sup>) | 45,073 | 0.75 | 0.72 | 3.3 | 4.3 | 2.2 |
| B (CD20<sup>+</sup>) | 13,624 | 0.68 | 0.64 | 6.6 | 7.5 | 5.8 |
| myeloid (CD68<sup>+</sup>) | 39,462 | 0.65 | 0.67 | −2.6 | −4.0 | 0.5 |

**Figure S3.**
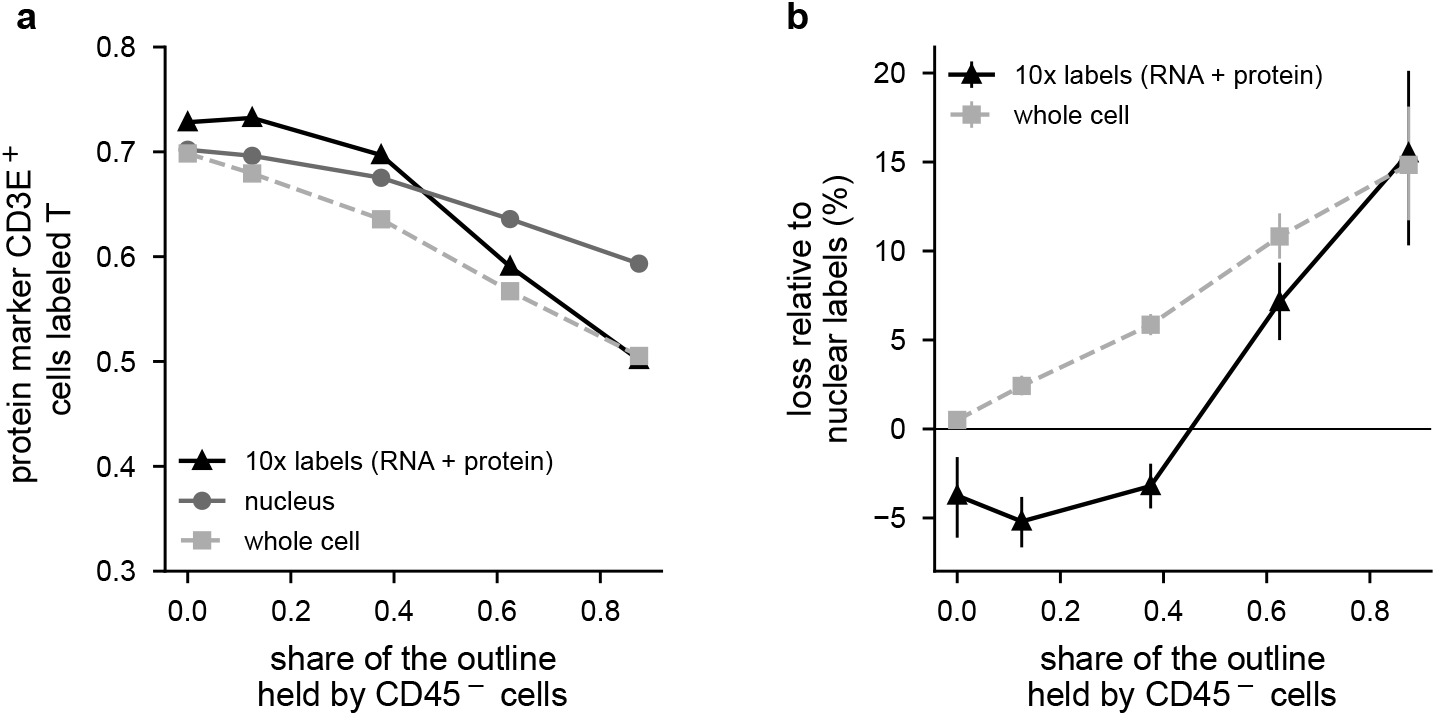
The vendor’s labels of renal cell carcinoma donor 2 against protein. On the 359,085 cells the published cell groups cover (52%), protein marker CD3E^+^ cells by the share of their outline held by CD45*^−^* cells. a, Share labeled T: by the published groups (RNA, and for immune and tumor cells RNA with protein, by weighted nearest neighbors), and by our labels from the whole nucleus and the whole cell. b, Loss relative to the nuclear labels, with 95% block intervals. Over all bins, the published groups recalled 0.74 of T cells defined by protein markers at precision 0.86.

**Table S8.**
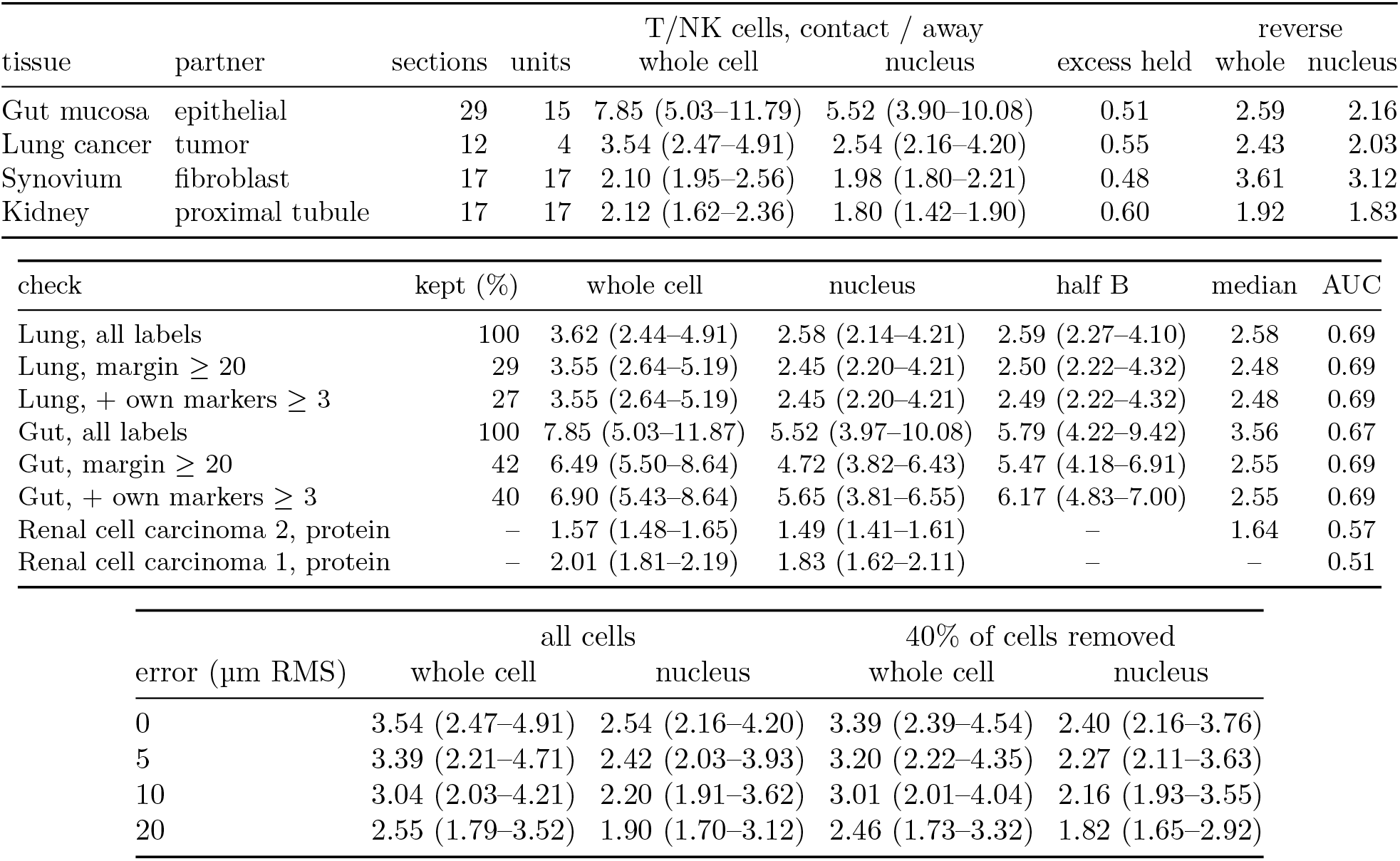
The neighbor’s RNA by contact. Partner-marker load (per 1000 counts) of T/NK cells at contact over away, medians over sections with 95% bootstrap intervals over donors or patients; excess held, median over sections of the contact-minus-away load in nuclear counts over that in whole-cell counts; reverse, the T/NK-marker load of partner cells at contact with T/NK cells over away. Middle block: checks for mislabeled cells, loads on the label genes; kept, the share of T/NK cells kept by the label margin (at least 20 nats) and own-marker counts (at least three); half B, the nuclear half the labels did not see; median, contact over away at the median nuclear load; AUC, the probability that a cell at contact carries a higher half B load than a cell away; renal cell carcinoma donor 2, T cells (CD3E^+^ CD45^+^ CD68*^−^* PanCK*^−^*) and PanCK^+^ tumor cells by protein, 213 tiles of 500 µm as units, 19 partner marker genes, AUC from nuclear counts; renal cell carcinoma donor 1, CD45-negative partners, 156 tiles, 1 partner marker gene. Lower block: lung, after an added positional error, with all cells and with 40% removed. Isolated nuclei (Slide-GoTags): T nuclei, tumor-marker load, 1.00 (0.78–1.08; above one in 7 of 15 sections); tumor nuclei, T-marker load, 1.28 (0.91–1.93; 9 sections).

| tissue | partner | sections | units | T/NK cells, contact / away |  |  | reverse |  |
|---|---|---|---|---|---|---|---|---|
|  |  |  |  | whole cell | nucleus | excess held | whole | nucleus |
| Gut mucosa | epithelial | 29 | 15 | 7.85 (5.03–11.79) | 5.52 (3.90–10.08) | 0.51 | 2.59 | 2.16 |
| Lung cancer | tumor | 12 | 4 | 3.54 (2.47–4.91) | 2.54 (2.16–4.20) | 0.55 | 2.43 | 2.03 |
| Synovium | fibroblast | 17 | 17 | 2.10 (1.95–2.56) | 1.98 (1.80–2.21) | 0.48 | 3.61 | 3.12 |
| Kidney | proximal tubule | 17 | 17 | 2.12 (1.62–2.36) | 1.80 (1.42–1.90) | 0.60 | 1.92 | 1.83 |
| check | kept (%) | whole cell | nucleus | half B | median | AUC |  |  |
| Lung, all labels | 100 | 3.62 (2.44–4.91) | 2.58 (2.14–4.21) | 2.59 (2.27–4.10) | 2.58 | 0.69 |  |  |
| Lung, margin ≥ 20 | 29 | 3.55 (2.64–5.19) | 2.45 (2.20–4.21) | 2.50 (2.22–4.32) | 2.48 | 0.69 |  |  |
| Lung, + own markers ≥ 3 | 27 | 3.55 (2.64–5.19) | 2.45 (2.20–4.21) | 2.49 (2.22–4.32) | 2.48 | 0.69 |  |  |
| Gut, all labels | 100 | 7.85 (5.03–11.87) | 5.52 (3.97–10.08) | 5.79 (4.22–9.42) | 3.56 | 0.67 |  |  |
| Gut, margin ≥ 20 | 42 | 6.49 (5.50–8.64) | 4.72 (3.82–6.43) | 5.47 (4.18–6.91) | 2.55 | 0.69 |  |  |
| Gut, + own markers ≥ 3 | 40 | 6.90 (5.43–8.64) | 5.65 (3.81–6.55) | 6.17 (4.83–7.00) | 2.55 | 0.69 |  |  |
| Renal cell carcinoma 2, protein | – | 1.57 (1.48–1.65) | 1.49 (1.41–1.61) | – | 1.64 | 0.57 |  |  |
| Renal cell carcinoma 1, protein | – | 2.01 (1.81–2.19) | 1.83 (1.62–2.11) | – | – | 0.51 |  |  |
|  | all cells |  | 40% of cells removed |  |  |  |  |  |
| error (µm RMS) | whole cell | nucleus | whole cell | nucleus |  |  |  |  |
| 0 | 3.54 (2.47–4.91) | 2.54 (2.16–4.20) | 3.39 (2.39–4.54) | 2.40 (2.16–3.76) |  |  |  |  |
| 5 | 3.39 (2.21–4.71) | 2.42 (2.03–3.93) | 3.20 (2.22–4.35) | 2.27 (2.11–3.63) |  |  |  |  |
| 10 | 3.04 (2.03–4.21) | 2.20 (1.91–3.62) | 3.01 (2.01–4.04) | 2.16 (1.93–3.55) |  |  |  |  |
| 20 | 2.55 (1.79–3.52) | 1.90 (1.70–3.12) | 2.46 (1.73–3.32) | 1.82 (1.65–2.92) |  |  |  |  |

**Table S9.**
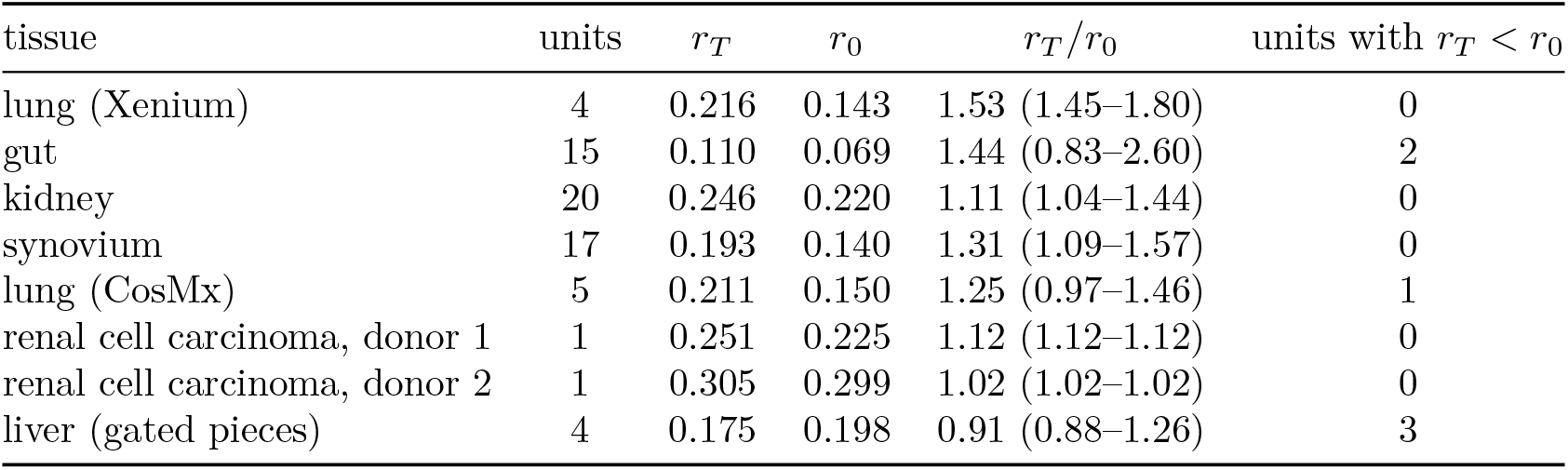
Change rates of test and null. Share of cells whose label changes from half A, in the whole-cell counts at the depth of half B (*r_T_*) and in half B (*r*_0_), per donor, patient, or piece (weighted by cells over its sections); medians over units, and the ratio *r_T_ /r*_0_ with its range. The extra-change ratio is undefined where *r_T_ < r*_0_. Per-unit values: Supplementary Data (rates_picked_units.csv).

| tissue | units | *r*<sub>*T*</sub> | *r*<sub>0</sub> | *r*<sub>*T*</sub>/*r*<sub>0</sub> | units with *r*<sub>*T*</sub> < *r*<sub>0</sub> |
|---|---|---|---|---|---|
| lung (Xenium) | 4 | 0.216 | 0.143 | 1.53 (1.45–1.80) | 0 |
| gut | 15 | 0.110 | 0.069 | 1.44 (0.83–2.60) | 2 |
| kidney | 20 | 0.246 | 0.220 | 1.11 (1.04–1.44) | 0 |
| synovium | 17 | 0.193 | 0.140 | 1.31 (1.09–1.57) | 0 |
| lung (CosMx) | 5 | 0.211 | 0.150 | 1.25 (0.97–1.46) | 1 |
| renal cell carcinoma, donor 1 | 1 | 0.251 | 0.225 | 1.12 (1.12–1.12) | 0 |
| renal cell carcinoma, donor 2 | 1 | 0.305 | 0.299 | 1.02 (1.02–1.02) | 0 |
| liver (gated pieces) | 4 | 0.175 | 0.198 | 0.91 (0.88–1.26) | 3 |

**Table S10.**
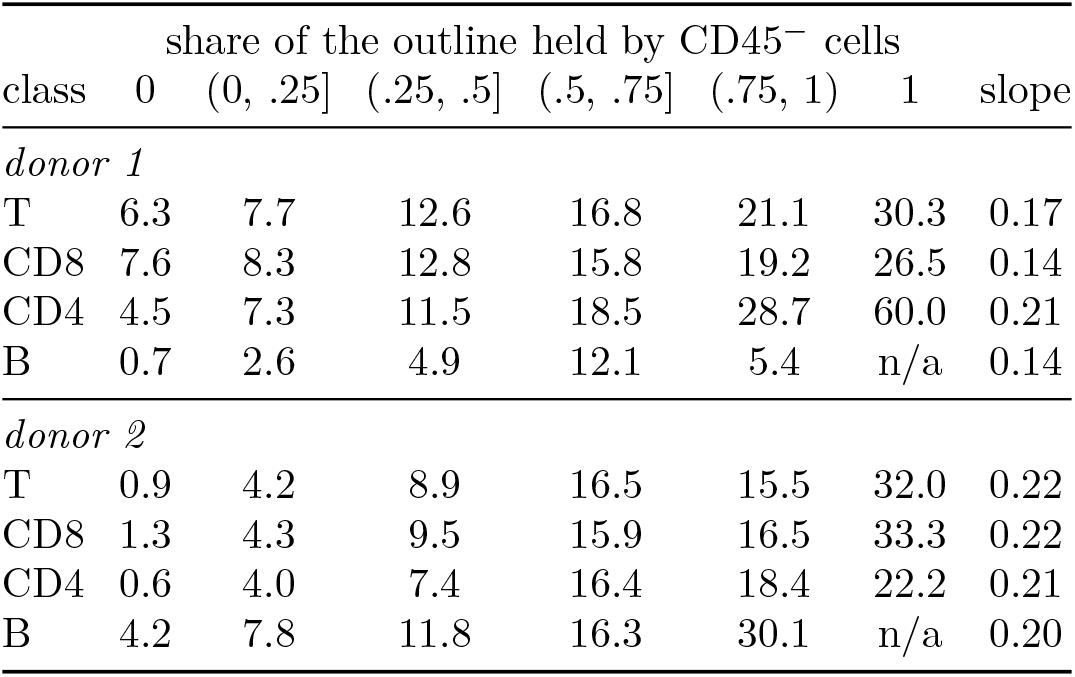
Renal cell carcinoma: protein loss by the non-immune share of the outline. Loss of the matching label (%, whole-cell counts against nuclear half B at equal depth) for protein-defined classes, by the share of the cell’s segmentation polygon held by CD45*^−^* cells (polygons densified every 1 µm; a point is held by the nearest point of another cell within 1 µm); slope, weighted least squares of the loss on the bin midpoints (bins of at least 200 cells). Share 1 holds few cells (328 and 254 T cells); 95% block intervals in the source tables. Cells with no neighbor within 1 µm (not shown) lost 11.8% (donor 1) and *−*0.1% (donor 2) of T labels. Candidates for the loss in donor 1 are neighbors above or below the section plane, cells within a few micrometers that the 1-µm polygon criterion counts as isolated, and differences between nuclear and cytoplasmic RNA; these data do not separate them. Within bins of the share of CD45-negative cells 30–100 µm away, the T-cell slopes were 0.15 (donor 1) and 0.21 (donor 2), compared with 0.17 and 0.22 over all cells.

**Figure S4.**
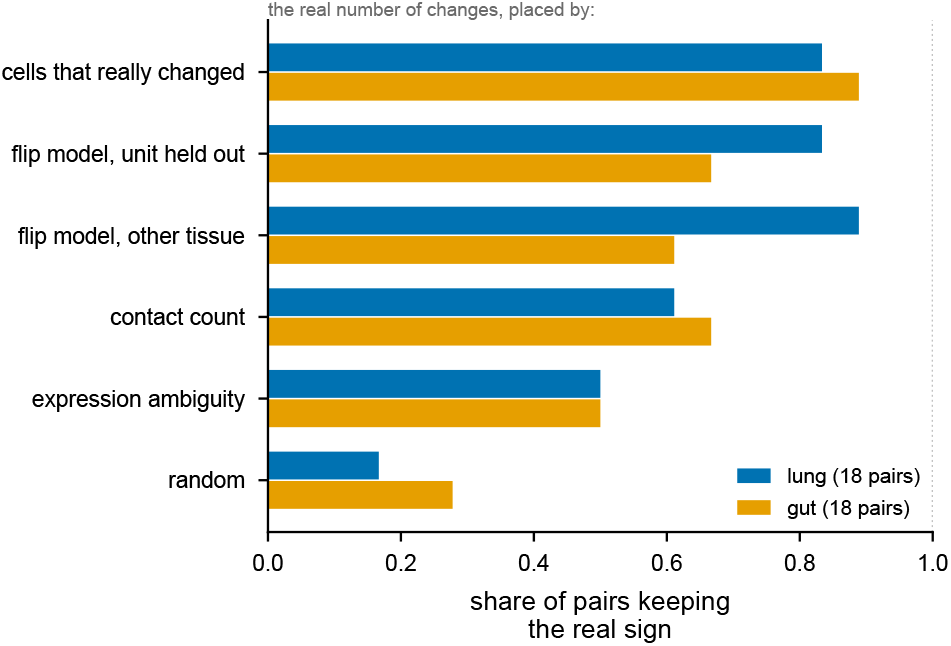
Placing errors where cells touch reproduces the real change. Share of type pairs whose real colocalization change (at least 0.02) keeps its sign when the real number of label changes is placed by each rule (lung donors, gut patients).

**Table S11.** Label loss by region. Loss of the immune label, whole-cell counts at the depth of nuclear half B against half B (%), for cells labeled as the immune type by half A, by the share of partner cells among the labeled cells within 25 µm (away, < 0.2; border, 0.2–0.5; inside, *≥* 0.5); medians over donors or patients (bins of at least 20 cells). Units: units with both bins whose loss is higher inside than away; difference, the median over units of each unit’s inside minus away (points), which need not equal the difference of the medians. Renal cell carcinomas: T cells defined by protein markers (CD3E^+^; strict, CD3E^+^ CD45^+^ CD68*^−^* PanCK*^−^*) and region cells typed by protein, loss pooled over the section, difference with a 95% tile-bootstrap interval. B cells near partner cells had uncertain half A labels (Results).

| cells, tissue | partner | away | border | inside | units | difference |
|---|---|---|---|---|---|---|
| *T/NK, labels* |  |  |  |  |  |  |
| gut, all masks | epithelium | 24 | 35 | 58 | 11 of 11 | +29 |
| gut, nucleus expansion | epithelium | 26 | 41 | 60 | — | — |
| gut, stain | epithelium | 18 | 24 | 38 | — | — |
| lung | tumor | 44 | 48 | 50 | 3 of 4 | +7 |
| synovium | fibroblasts | 20 | 32 | 38 | 15 of 16 | +16 |
| kidney | loop of Henle | 29 | 25 | 18 | 2 of 8 | −5 |
| kidney | proximal tubule | 28 | 32 | 26 | 3 of 7 | −1 |
| kidney | non-immune | 19 | 27 | 29 | 7 of 9 | +6 |
| *T, protein* |  |  |  |  |  |  |
| renal cell carcinoma 1 | CD45<sup>−</sup> | 7.3 | 10.5 | 14.3 | — | +7.0 (5.6 to 8.4) |
| strict T | CD45<sup>−</sup> | 6.7 | 9.7 | 13.7 | — | +7.0 (5.6 to 8.4) |
| renal cell carcinoma 2 | CD45<sup>−</sup> | 1.8 | 5.0 | 6.6 | — | +4.9 (3.7 to 6.0) |
| strict T | CD45<sup>−</sup> | 1.1 | 4.4 | 6.0 | — | +4.9 (3.8 to 6.1) |
| renal cell carcinoma 2 | PanCK<sup>+</sup> | 4.3 | 8.0 | 12.4 | — | +8.1 (2.9 to 13.1) |
| strict T | PanCK<sup>+</sup> | 3.8 | 6.0 | 6.9 | — | +3.1 (−4.0 to 9.5) |
| *B and plasma cells, labels* |  |  |  |  |  |  |
| B, gut | epithelium | 9 | 19 | 21 | 5 of 6 | — |
| B, lung | tumor | 24 | 38 | 41 | 4 of 4 | — |
| B, synovium | fibroblasts | 20 | 30 | 50 | 8 of 11 | — |
| plasma, gut | epithelium | 12 | 15 | 18 | 6 of 10 | — |
| plasma, synovium | fibroblasts | 5 | 7 | 15 | 10 of 13 | — |

**Table S12.** Which T/NK calls lose their labels, matched on nuclear counts. Loss of calls positive for a gene set (at least one nuclear count) minus that of negative calls (points), all regions together; matched, within five strata of nuclear counts per unit (Mantel–Haenszel weights), median over units, with the crude difference in parentheses and the units with a positive matched difference over all units. Last row: inside minus away, matched in the same way. Partners as in Supplementary Table S11 (kidney, the proximal tubule). Gene sets in Supplementary Methods.

| set | gut | lung | synovium | kidney |
|---|---|---|---|---|
| cytotoxic | −0.2 (+3.5); 6/13 | +1.2 (+2.5); 3/4 | −0.8 (−2.9); 8/17 | −3.7 (−3.1); 6/17 |
| CD8 | +3.6 (+2.1); 8/12 | −5.4 (−5.5); 0/4 | −1.2 (−3.4); 2/16 | −0.8 (−1.9); 8/17 |
| CD4 | −6.2 (−7.7); 3/11 | +2.4 (+3.2); 3/4 | −0.4 (−3.2); 6/16 | −5.7 (−3.2); 4/14 |
| *γδ* | +20.1 (+18.6); 2/2 | +7.8 (+8.1); 4/4 | −0.8 (−4.0); 5/16 | −0.5 (+1.0); 1/2 |
| tissue-resident | +10.7 (+3.5); 3/4 | +0.1 (+0.6); 2/4 | −0.5 (−3.3); 7/16 | −2.2 (+4.1); 6/13 |
| naive-like | −8.7 (−12.5); 1/10 | +1.5 (+2.4); 2/4 | −5.5 (−8.4); 0/16 | −2.3 (−3.6); 3/9 |
| proliferating | +2.2 (+1.0); 8/11 | −5.5 (−6.4); 0/4 | −1.2 (−2.5); 5/16 | −9.6 (−9.3); 0/3 |
| activated | −0.9 (−2.4); 5/12 | +11.6 (+10.5); 4/4 | −2.9 (−5.8); 2/16 | −0.7 (−1.2); 6/13 |
| exhausted | +4.0 (−3.2); 7/13 | +1.3 (+2.0); 3/4 | −0.9 (−4.4); 2/16 | −1.4 (−0.5); 7/15 |
| inside vs away | +26.0 (+29.3); 11/11 | +6.6 (+7.0); 3/4 | +14.1 (+14.9); 15/17 | +5.3 (+1.5); 4/6 |

**Table S13.** Who the T/NK calls inside the partner region are. T/NK calls of nuclear half A by region (as in Supplementary Table S11) and, for comparison, the partner cells themselves; medians per donor or patient, then over them. Areas from the cell tables (µm^2^); counts, nuclear and whole-cell; markers, nuclear counts of T-cell, partner, and cytotoxic genes per 1000 nuclear counts (Supplementary Methods); runner-up, share of calls whose second-best half A label is the partner; half B, share of calls that nuclear half B also labels T/NK.

|  | area |  | counts |  | markers per 1000 |  |  |  |  |
|---|---|---|---|---|---|---|---|---|---|
|  | nucleus | cell | nucleus | whole | T cell | partner | cytotoxic | runner-up | half B |
| *gut, partner epithelium* |  |  |  |  |  |  |  |  |  |
| T/NK away | 28.6 | 52 | 30 | 52 | 282 | 0 | 0 | 0.00 | 0.67 |
| T/NK border | 28.2 | 65 | 28 | 49 | 229 | 0 | 0 | 0.03 | 0.69 |
| T/NK inside | 27.3 | 83 | 30 | 57 | 190 | 0 | 48 | 0.22 | 0.68 |
| epithelial cells | 29.9 | 84 | 38 | 77 | 0 | 96 | 0 | — | — |
| *lung, partner tumor* |  |  |  |  |  |  |  |  |  |
| T/NK away | 25.9 | 68 | 40 | 84 | 91 | 0 | 0 | 0.01 | 0.74 |
| T/NK border | 26.5 | 82 | 41 | 94 | 73 | 0 | 5 | 0.07 | 0.75 |
| T/NK inside | 26.6 | 89 | 43 | 104 | 69 | 19 | 24 | 0.24 | 0.75 |
| tumor cells | 50.3 | 135 | 82 | 200 | 0 | 112 | 0 | — | — |

**Table S14.**
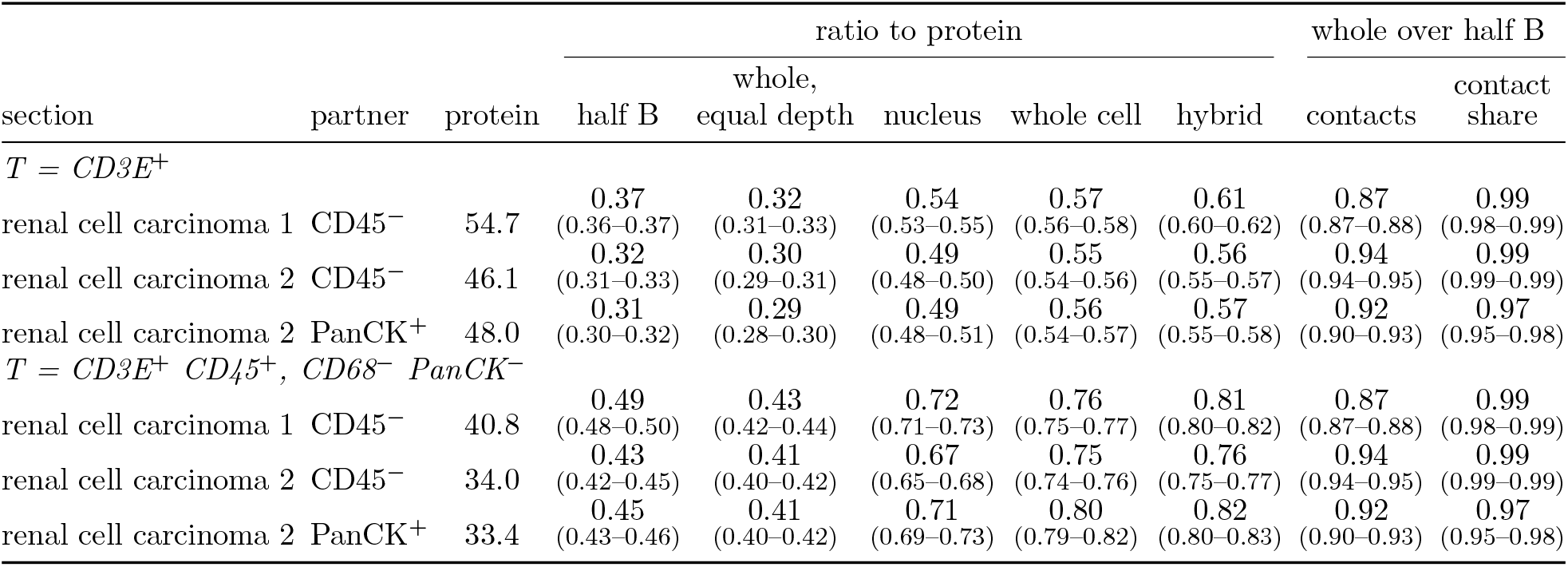
Renal cell carcinoma: T-cell contacts scored against protein. T cells contacting a partner cell (six nearest within 30 µm) per 100 partner cells, partners typed by protein (CD45-negative, or PanCK-positive; pan-cytokeratin does not separate in donor 1). Protein markers: the count with T cells called by protein markers. Ratio to protein: the same count with T cells called by each labeling, over the protein value; half B and whole (equal depth) at the depth of nuclear half B; hybrid, the nuclear label where the nucleus held at least ten counts, else the whole-cell label; cells left unlabeled count as non-T. Whole over half B, equal depth: the count (contacts) and the share of T cells contacting a partner (contact share). 95% tile-bootstrap intervals in brackets, paired between the two count sets of a ratio. The positional part of the fall in contacts (Results) was *−*2.6 and *−*2.0 points with partners typed by protein markers, against *−*6.1 and *−*4.4 with the label-only check.

| section | partner | protein | ratio to protein |  |  |  |  | whole over half B |  |
|---|---|---|---|---|---|---|---|---|---|
| section | partner | protein | half B | whole,<br>equal depth | nucleus | whole cell | hybrid | contacts | contact<br>share |
| *T* = *CD3E*<sup>+</sup> |  |  |  |  |  |  |  |  |  |
| renal cell carcinoma 1 | CD45<sup>−</sup> | 54.7 | 0.37<br>(0.36–0.37) | 0.32<br>(0.31–0.33) | 0.54<br>(0.53–0.55) | 0.57<br>(0.56–0.58) | 0.61<br>(0.60–0.62) | 0.87<br>(0.87–0.88) | 0.99<br>(0.98–0.99) |
| renal cell carcinoma 2 | CD45<sup>−</sup> | 46.1 | 0.32<br>(0.31–0.33) | 0.30<br>(0.29–0.31) | 0.49<br>(0.48–0.50) | 0.55<br>(0.54–0.56) | 0.56<br>(0.55–0.57) | 0.94<br>(0.94–0.95) | 0.99<br>(0.99–0.99) |
| renal cell carcinoma 2 | PanCK<sup>+</sup> | 48.0 | 0.31<br>(0.30–0.32) | 0.29<br>(0.28–0.30) | 0.49<br>(0.48–0.51) | 0.56<br>(0.54–0.57) | 0.57<br>(0.55–0.58) | 0.92<br>(0.90–0.93) | 0.97<br>(0.95–0.98) |
| *T* = *CD3E*<sup>+</sup> *CD45*<sup>+</sup>, *CD68*<sup>−</sup> *PanCK*<sup>−</sup> |  |  |  |  |  |  |  |  |  |
| renal cell carcinoma 1 | CD45<sup>−</sup> | 40.8 | 0.49<br>(0.48–0.50) | 0.43<br>(0.42–0.44) | 0.72<br>(0.71–0.73) | 0.76<br>(0.75–0.77) | 0.81<br>(0.80–0.82) | 0.87<br>(0.87–0.88) | 0.99<br>(0.98–0.99) |
| renal cell carcinoma 2 | CD45<sup>−</sup> | 34.0 | 0.43<br>(0.42–0.45) | 0.41<br>(0.40–0.42) | 0.67<br>(0.65–0.68) | 0.75<br>(0.74–0.76) | 0.76<br>(0.75–0.77) | 0.94<br>(0.94–0.95) | 0.99<br>(0.99–0.99) |
| renal cell carcinoma 2 | PanCK<sup>+</sup> | 33.4 | 0.45<br>(0.43–0.46) | 0.41<br>(0.40–0.42) | 0.71<br>(0.69–0.73) | 0.80<br>(0.79–0.82) | 0.82<br>(0.80–0.83) | 0.92<br>(0.90–0.93) | 0.97<br>(0.95–0.98) |

**Table S15.** Renal cell carcinoma: the remedy scored by protein. Two donors. Recall: share of protein-defined cells given the matching label, per count set; precision: share of cells given the label that are protein-positive for the class. Half B, nuclear half B; whole, equal depth, whole-cell counts minus half A thinned to the depth of B; B + outside, half B plus all outside counts; N-aware, the same counts with the neighbor-aware likelihood; nucleus and whole cell, full counts; whole, N-aware, the whole nucleus plus the outside counts with the neighbor-aware likelihood. Weight sensitivity (Methods): with contact alone, and with the regional and ambient weights halved or doubled, the likelihood closed 0.06–0.48 of the gap in T recall between whole cell and nucleus, least with contact alone, and in all 8 runs it remained below the nucleus and lowered myeloid recall.

| protein class | recall |  |  |  |  |  |  | precision |  |  |
|---|---|---|---|---|---|---|---|---|---|---|
|  | half<br>B | whole,<br>equal<br>depth | B +<br>outside | B +<br>outside,<br>N-aware | nucleus | whole<br>cell | whole,<br>N-aware | half<br>B | B +<br>outside | B +<br>outside,<br>N-aware |
| *donor 1* |  |  |  |  |  |  |  |  |  |  |
| T (CD3E<sup>+</sup>) | 0.63 | 0.57 | 0.54 | 0.58 | 0.64 | 0.58 | 0.61 | 0.92 | 0.92 | 0.91 |
| CD8 (CD8A<sup>+</sup>) | 0.74 | 0.66 | 0.63 | 0.68 | 0.74 | 0.67 | 0.71 | 0.63 | 0.63 | 0.63 |
| CD4 (CD4<sup>+</sup>CD3E<sup>+</sup>) | 0.77 | 0.71 | 0.69 | 0.71 | 0.78 | 0.73 | 0.75 | 0.22 | 0.23 | 0.22 |
| B (CD20<sup>+</sup>) | 0.87 | 0.86 | 0.85 | 0.82 | 0.87 | 0.86 | 0.84 | 0.82 | 0.83 | 0.80 |
| myeloid (CD68<sup>+</sup>) | 0.82 | 0.85 | 0.86 | 0.81 | 0.81 | 0.85 | 0.81 | 0.53 | 0.51 | 0.53 |
| *donor 2* |  |  |  |  |  |  |  |  |  |  |
| T (CD3E<sup>+</sup>) | 0.61 | 0.59 | 0.58 | 0.59 | 0.63 | 0.61 | 0.61 | 0.92 | 0.93 | 0.92 |
| CD8 (CD8A<sup>+</sup>) | 0.73 | 0.69 | 0.69 | 0.70 | 0.74 | 0.71 | 0.72 | 0.47 | 0.48 | 0.47 |
| CD4 (CD4<sup>+</sup>CD3E<sup>+</sup>) | 0.75 | 0.72 | 0.72 | 0.73 | 0.76 | 0.74 | 0.74 | 0.46 | 0.46 | 0.45 |
| B (CD20<sup>+</sup>) | 0.68 | 0.64 | 0.61 | 0.63 | 0.69 | 0.65 | 0.66 | 0.68 | 0.73 | 0.68 |
| myeloid (CD68<sup>+</sup>) | 0.65 | 0.67 | 0.67 | 0.65 | 0.65 | 0.66 | 0.65 | 0.44 | 0.46 | 0.46 |

**Table S16.** Labeling from the nucleus where it holds enough counts. Recall of protein-positive cells, with every protein-positive cell counted (unlabeled cells as missed). Labeled: share of protein-positive cells with a nuclear label (at least ten nuclear counts). Hybrid: the nuclear label where the nucleus held at least ten counts, else the whole-cell label. Lung, CD8^+^ cells by immunohistochemistry, labeled T/NK (deposited masks); renal cell carcinoma, CD3E^+^ cells labeled T; uterine cancer, CD45RO^+^ cells labeled T. Lower block, renal cell carcinoma: precision of the T calls (share of cells labeled T that are CD3E^+^) and recall of CD68^+^ cells labeled myeloid, all cells counted.

| section | labeled from the nucleus | nucleus | whole cell | hybrid |
|---|---|---|---|---|
| lung L1 | 0.78 | 0.36 | 0.31 | 0.39 |
| lung L3 | 0.90 | 0.65 | 0.42 | 0.66 |
| renal cell carcinoma 1 | 0.79 | 0.51 | 0.54 | 0.56 |
| renal cell carcinoma 2 | 0.71 | 0.45 | 0.51 | 0.51 |
| uterine cancer, MERSCOPE | 0.76 | 0.28 | 0.20 | 0.30 |
| renal cell carcinoma 1, T precision |  | 0.91 | 0.92 | 0.91 |
| renal cell carcinoma 2, T precision |  | 0.92 | 0.93 | 0.92 |
| renal cell carcinoma 1, myeloid recall |  | 0.64 | 0.82 | 0.79 |
| renal cell carcinoma 2, myeloid recall |  | 0.46 | 0.56 | 0.54 |

**Table S17.**
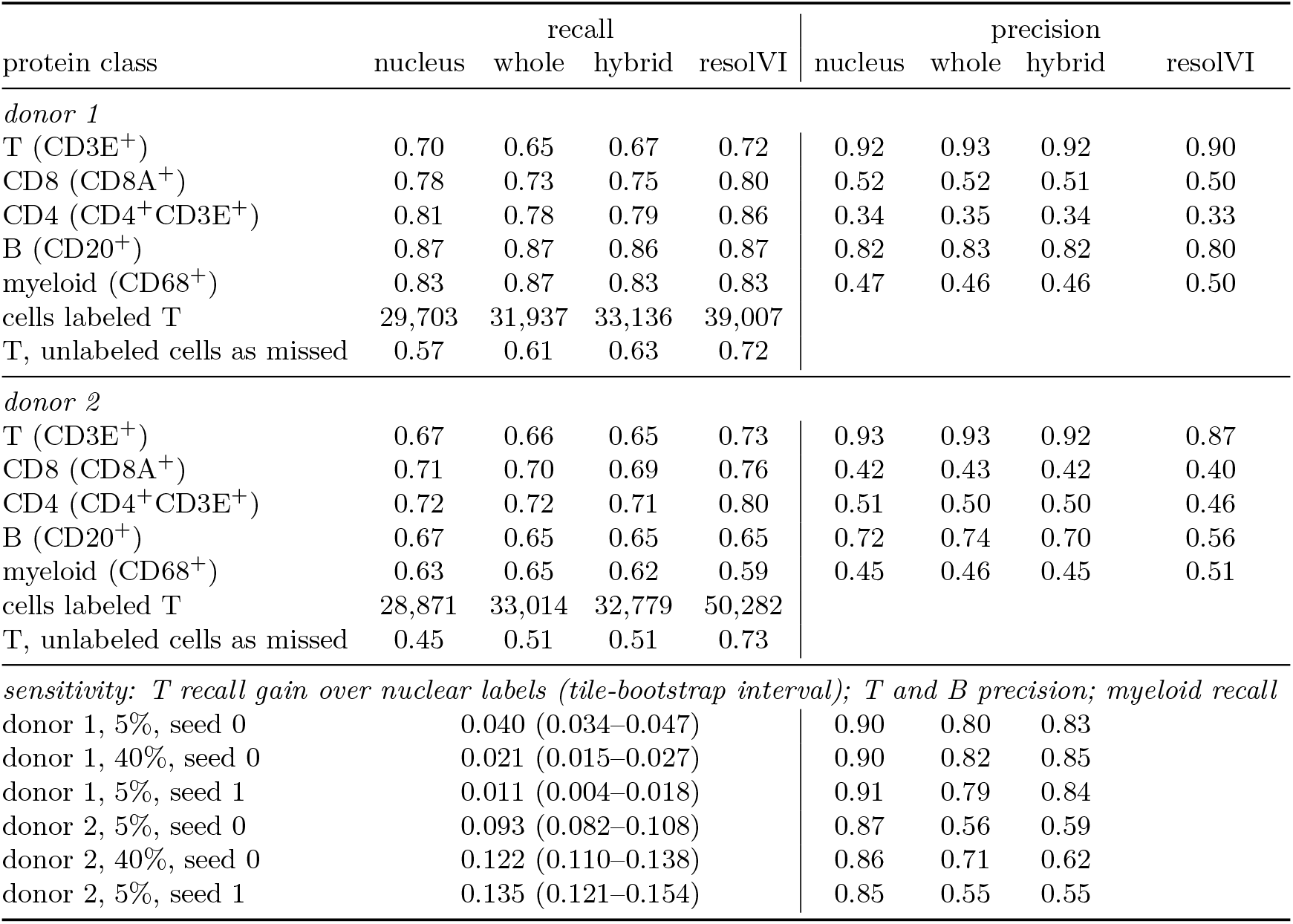
Renal cell carcinoma: resolVI relabeling scored by protein. The three windows per donor with the most T/NK cells (120,619 and 137,307 cells); every labeling scored on the same cells. Recall and precision as in Supplementary Table S15; nucleus and whole, our labels from full nuclear and whole-cell counts; hybrid, the nuclear label where the nucleus labeled the cell, else the whole-cell label (Supplementary Table S16); resolVI, semi-supervised resolVI started from the whole-cell labels (Methods). Recall counts the cells each labeling labeled; our classifier leaves cells with too few counts unlabeled (T cells labeled: nucleus 0.82 and 0.67, whole cell and hybrid 0.94 and 0.78), whereas resolVI labels every cell, so the row with unlabeled cells counted as missed compares all four on the same footing. The windows were rebuilt from the resolVI settings and reproduced the nuclear and whole-cell recalls of that run. Lower block: the first run (5% uncertainty filter, seed 0), the authors’ filter for fine labels (40%), and a second seed; columns there are the gain in T recall over nuclear labels, T precision, B precision, and myeloid recall. Paired differences in T recall, resolVI minus nucleus, first run: 0.040 (0.034–0.047) and 0.093 (0.082–0.108), tile bootstrap. Starting labels kept for training, 0.25–0.38 and 0.05–0.22 of cells per window; resolVI kept the starting label of 0.86–0.89 and 0.71–0.86. In every run, resolVI recalled more T cells defined by protein markers than nuclear labels (by 0.01–0.04 in donor 1 and 0.09–0.14 in donor 2), and in donor 1 it reduced the extra loss at contact with CD68^+^ cells, by calling more cells T, so T precision fell in every run.

| protein class | recall |  |  |  | precision |  |  |  |
|---|---|---|---|---|---|---|---|---|
|  | nucleus | whole | hybrid | resolVI | nucleus | whole | hybrid | resolVI |
| *donor 1* |  |  |  |  |  |  |  |  |
| T (CD3E<sup>+</sup>) | 0.70 | 0.65 | 0.67 | 0.72 | 0.92 | 0.93 | 0.92 | 0.90 |
| CD8 (CD8A<sup>+</sup>) | 0.78 | 0.73 | 0.75 | 0.80 | 0.52 | 0.52 | 0.51 | 0.50 |
| CD4 (CD4<sup>+</sup>CD3E<sup>+</sup>) | 0.81 | 0.78 | 0.79 | 0.86 | 0.34 | 0.35 | 0.34 | 0.33 |
| B (CD20<sup>+</sup>) | 0.87 | 0.87 | 0.86 | 0.87 | 0.82 | 0.83 | 0.82 | 0.80 |
| myeloid (CD68<sup>+</sup>) | 0.83 | 0.87 | 0.83 | 0.83 | 0.47 | 0.46 | 0.46 | 0.50 |
| cells labeled T | 29,703 | 31,937 | 33,136 | 39,007 |  |  |  |  |
| T, unlabeled cells as missed | 0.57 | 0.61 | 0.63 | 0.72 |  |  |  |  |
| *donor 2* |  |  |  |  |  |  |  |  |
| T (CD3E<sup>+</sup>) | 0.67 | 0.66 | 0.65 | 0.73 | 0.93 | 0.93 | 0.92 | 0.87 |
| CD8 (CD8A<sup>+</sup>) | 0.71 | 0.70 | 0.69 | 0.76 | 0.42 | 0.43 | 0.42 | 0.40 |
| CD4 (CD4<sup>+</sup>CD3E<sup>+</sup>) | 0.72 | 0.72 | 0.71 | 0.80 | 0.51 | 0.50 | 0.50 | 0.46 |
| B (CD20<sup>+</sup>) | 0.67 | 0.65 | 0.65 | 0.65 | 0.72 | 0.74 | 0.70 | 0.56 |
| myeloid (CD68<sup>+</sup>) | 0.63 | 0.65 | 0.62 | 0.59 | 0.45 | 0.46 | 0.45 | 0.51 |
| cells labeled T | 28,871 | 33,014 | 32,779 | 50,282 |  |  |  |  |
| T, unlabeled cells as missed | 0.45 | 0.51 | 0.51 | 0.73 |  |  |  |  |
| *sensitivity: T recall gain over nuclear labels (tile-bootstrap interval); T and B precision; myeloid recall* |  |  |  |  |  |  |  |  |
| donor 1, 5%, seed 0 | 0.040 (0.034–0.047) |  |  |  | 0.90 | 0.80 | 0.83 |  |
| donor 1, 40%, seed 0 | 0.021 (0.015–0.027) |  |  |  | 0.90 | 0.82 | 0.85 |  |
| donor 1, 5%, seed 1 | 0.011 (0.004–0.018) |  |  |  | 0.91 | 0.79 | 0.84 |  |
| donor 2, 5%, seed 0 | 0.093 (0.082–0.108) |  |  |  | 0.87 | 0.56 | 0.59 |  |
| donor 2, 40%, seed 0 | 0.122 (0.110–0.138) |  |  |  | 0.86 | 0.71 | 0.62 |  |
| donor 2, 5%, seed 1 | 0.135 (0.121–0.154) |  |  |  | 0.85 | 0.55 | 0.55 |  |

**Table S18.**
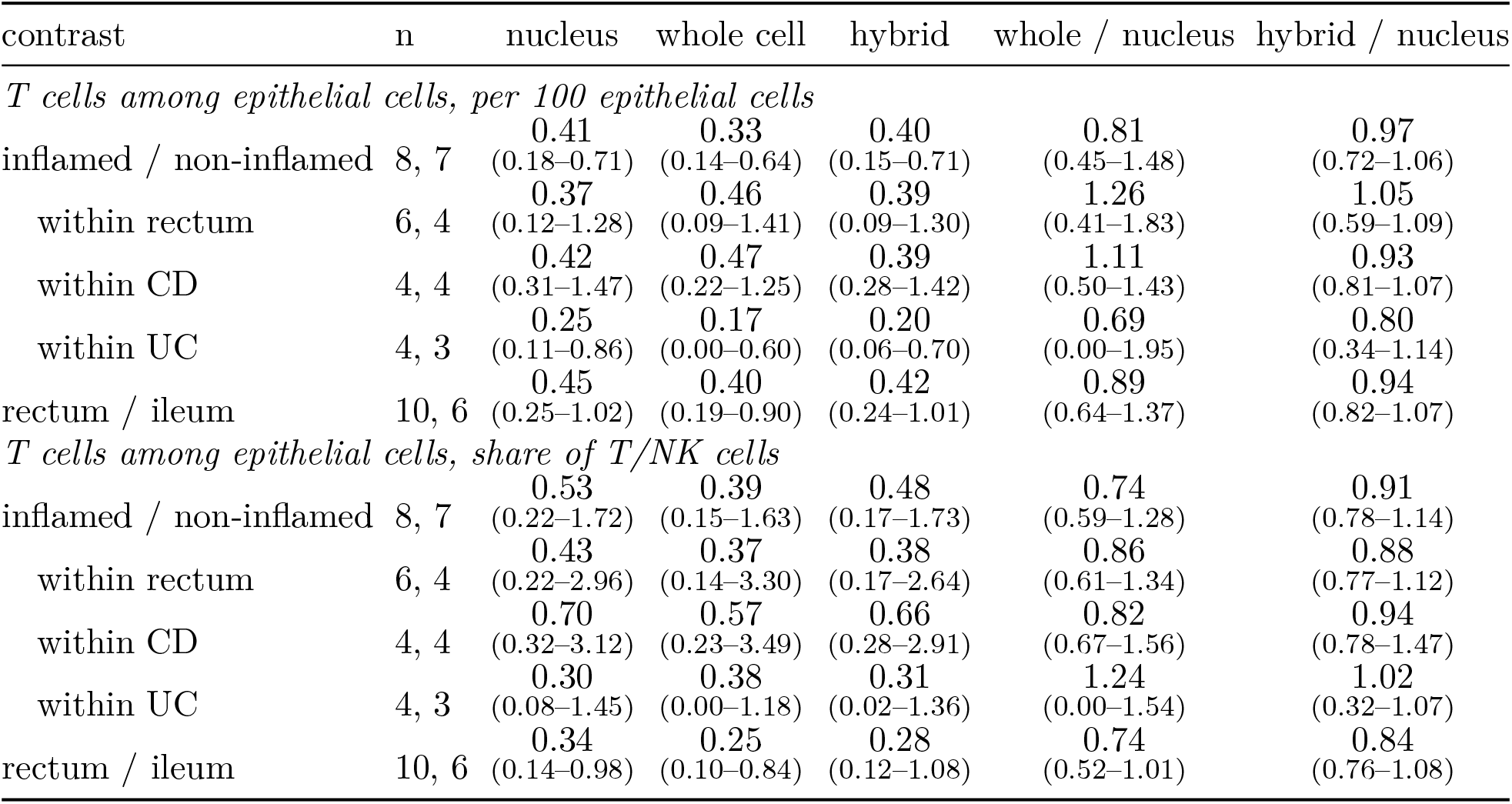
T cells among epithelial cells by condition and labeling, gut. The 16 sections on the 377-gene panel, 11 patients; conditions from the GEO sample titles. T cells among epithelial cells: T/NK cells with at least half epithelial cells among the labeled cells within 25 µm, by the labeling named. Ratio of the medians over patients between the two levels, with 95% bootstrap intervals over patients (2000 draws, the same for every labeling); n, patient groups in each level (a patient with sections in both levels counts in both). Whole / nucleus and hybrid / nucleus, the ratio of the contrast between labelings: one if the labeling does not change it. The share of T/NK cells among epithelial cells does not depend on how many epithelial cells a labeling finds; whole-cell labels find more, which lowers the count per 100 epithelial cells. Because some patients count in both levels, the intervals are approximate; n/a, an interval end that the bootstrap could not estimate, because a resampled level had a median of zero T cells among epithelial cells.

| contrast | n | nucleus | whole cell | hybrid | whole / nucleus | hybrid / nucleus |
|---|---|---|---|---|---|---|
| *T cells among epithelial cells, per 100 epithelial cells* |  |  |  |  |  |  |
| inflamed / non-inflamed | 8, 7 | 0.41<br>(0.18–0.71) | 0.33<br>(0.14–0.64) | 0.40<br>(0.15–0.71) | 0.81<br>(0.45–1.48) | 0.97<br>(0.72–1.06) |
| within rectum | 6, 4 | 0.37<br>(0.12–1.28) | 0.46<br>(0.09–1.41) | 0.39<br>(0.09–1.30) | 1.26<br>(0.41–1.83) | 1.05<br>(0.59–1.09) |
| within CD | 4, 4 | 0.42<br>(0.31–1.47) | 0.47<br>(0.22–1.25) | 0.39<br>(0.28–1.42) | 1.11<br>(0.50–1.43) | 0.93<br>(0.81–1.07) |
| within UC | 4, 3 | 0.25<br>(0.11–0.86) | 0.17<br>(0.00–0.60) | 0.20<br>(0.06–0.70) | 0.69<br>(0.00–1.95) | 0.80<br>(0.34–1.14) |
| rectum / ileum | 10, 6 | 0.45<br>(0.25–1.02) | 0.40<br>(0.19–0.90) | 0.42<br>(0.24–1.01) | 0.89<br>(0.64–1.37) | 0.94<br>(0.82–1.07) |
| *T cells among epithelial cells, share of T/NK cells* |  |  |  |  |  |  |
| inflamed / non-inflamed | 8, 7 | 0.53<br>(0.22–1.72) | 0.39<br>(0.15–1.63) | 0.48<br>(0.17–1.73) | 0.74<br>(0.59–1.28) | 0.91<br>(0.78–1.14) |
| within rectum | 6, 4 | 0.43<br>(0.22–2.96) | 0.37<br>(0.14–3.30) | 0.38<br>(0.17–2.64) | 0.86<br>(0.61–1.34) | 0.88<br>(0.77–1.12) |
| within CD | 4, 4 | 0.70<br>(0.32–3.12) | 0.57<br>(0.23–3.49) | 0.66<br>(0.28–2.91) | 0.82<br>(0.67–1.56) | 0.94<br>(0.78–1.47) |
| within UC | 4, 3 | 0.30<br>(0.08–1.45) | 0.38<br>(0.00–1.18) | 0.31<br>(0.02–1.36) | 1.24<br>(0.00–1.54) | 1.02<br>(0.32–1.07) |
| rectum / ileum | 10, 6 | 0.34<br>(0.14–0.98) | 0.25<br>(0.10–0.84) | 0.28<br>(0.12–1.08) | 0.74<br>(0.52–1.01) | 0.84<br>(0.76–1.08) |

**Figure S5.**
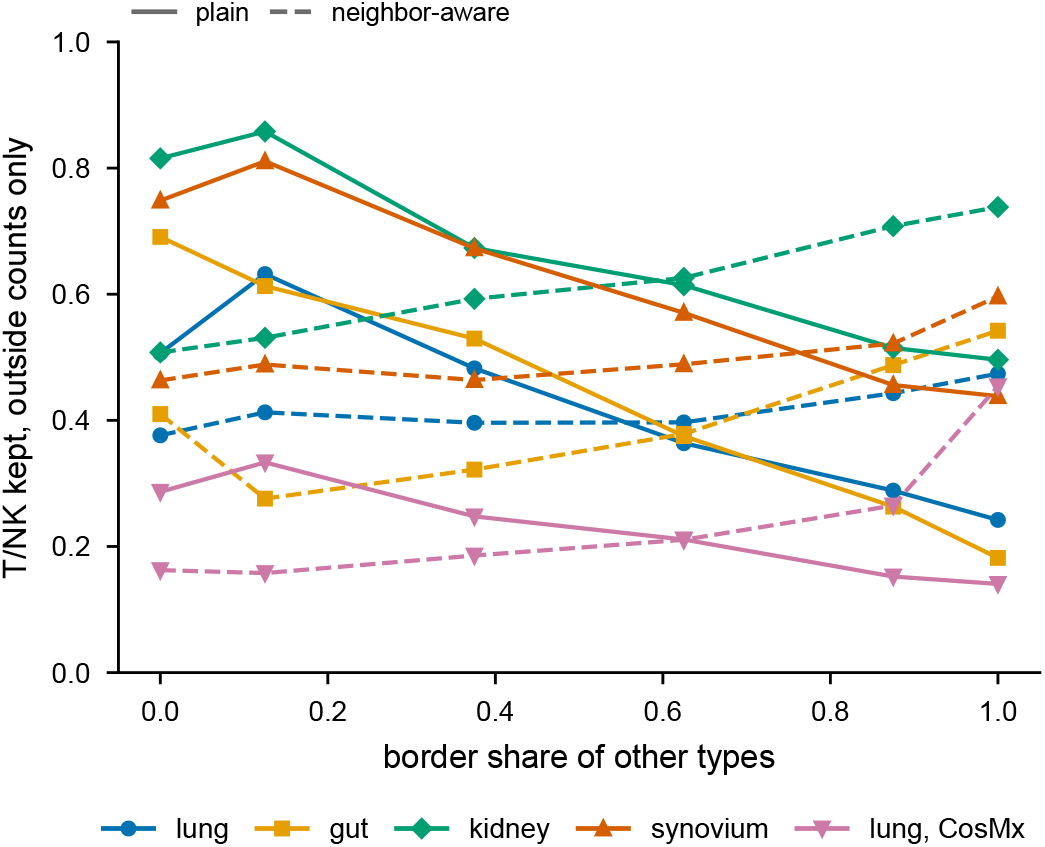
T/NK kept from outside counts against border share, plain and neighbor-aware. T/NK labels kept by the counts outside the nucleus over nuclear half B, against the share of the border held by other types, plain (solid) and with the neighbor-aware likelihood (dashed); medians over donors or patients; border share from the nuclear-disk proxy (Methods). The slopes are summarized in Fig. 6c.

#### Pieces and units

The four CosMx slides of GSE312698 hold ten samples; gaps between fields of view gave 15 pieces of at least 2000 cells (549,602 cells). In 4 pieces the median cell had no nuclear transcript, so the test cannot be run on them; in the others, fields of view with fewer than a quarter of transcripts in the nucleus were removed (499,417 cells remain). One slide holds two large sections, presumably the two explants; the other three hold needle-biopsy strips, some broken into several pieces, and the mapping of pieces to patients is not deposited, so we report pieces rather than patients.

#### Labels

Table S19 lists the classifier settings tried. With the per-gene platform factor (fitted from hard labels and refitted five times) or a class prior fitted by expectation maximization, the labels of the biopsy pieces on two slides collapsed onto cholangiocyte (nuclear) or endothelial (whole cell). The agreement of the nuclear halves did not flag this: it was high because both halves received the same incorrect class. Without the platform factor nothing collapsed, and the halves of the same pieces agreed in only 0.30–0.49 of cells. A reference-free call (for each cell, the class whose canonical panel markers take the largest share of its counts relative to the tissue average) agreed with nuclear labels in Leuven 1 in 0.65–0.75 of cells with the chosen setting, compared with 0.59–0.65 with the default; in the other pieces, the marker call itself was nearly uniform across classes, consistent with weak signal (the panel holds only two of twelve canonical hepatocyte markers). In practice, the choice of setting, therefore, rests on the two explant sections. The reference has normal donors only and is dominated by dissociated cells, in which hepatocytes are under-represented; a nuclear reference or the authors’ annotation would be better.

#### Test

On the 4 pieces that passed, with the setting used, one piece was evaluable for the extra-change ratio (1.48); with 30 markers and a fitted prior, 2 pieces (1.50–1.53, intervals above one in 2). The default labels collapse and are not used. Random relabeling of 10% of the whole-cell labels raised contacts per 100 hepatocytes (T/NK, B, and myeloid), which the whole-cell labels had lowered, in 12 of 12 piece–type pairs, as in the Xenium tissues.

#### Stains

Spearman correlation of each CosMx stain with the cell’s own marker transcripts against its nearest neighbor’s (medians over pieces): CD45 with PTPRC 0.04 compared with 0.02, CD68 with CD68 0.05 compared with 0.02, pan-cytokeratin with KRT8/18/19/7 0.05 compared with 0.02. As with CD3 in CosMx lung, the stains do not describe the stained cell well enough to serve as truth. In a third CosMx dataset, three tissue microarrays of head and neck squamous cell carcinoma [47], the CD3 stain did not separate positive cells (Ashman *D* 0.74–0.94) and correlated with the cell’s own CD3 transcripts (up to 0.19) barely more than with its nearest neighbor’s (up to 0.14); pan-cytokeratin separated, but followed the neighbor’s keratin transcripts almost as closely as the cell’s own (0.52–0.55 compared with 0.56–0.59; hnscc_stain_check.py). A fourth CosMx dataset, a breast cancer section with the 64-plex immuno-oncology protein panel and the whole-transcriptome panel on the same slide and one segmentation [46] (150,142 cells), carries real antibodies rather than morphology stains. CD3-high cells there carried 2.8 times more T-cell signature transcripts than other cells, but their nearest neighbors carried 2.1 times more; for CD3, CD8, and CD4, the ratio of the two folds was at most 1.36, compared with 2.1–4.6 for Xenium Protein (Supplementary Methods; cosmx_protein_check.py). The CosMx antibody signal marks T-rich regions more than individual T cells, so this dataset cannot serve as same-cell truth either.

**Table S19.** Liver: classifier settings. Largest share of one class in any piece and count set (nuclear or whole cell); pieces in which a non-hepatocyte class exceeds 60%; agreement of the nuclear halves (range over pieces) for the explant sections (Leuven 1), the biopsies of slide 3, and those of slides 2 and 4; median agreement of nuclear half B with the reference-free marker call. All settings exclude the 207 over-detected probes. *^∗^*Used for the liver test.

| setting | largest (%) | collapsed | halves L1 | halves L3 | halves L2, L4 | markers L1/other |
|---|---|---|---|---|---|---|
| platform factor (default) | 94 | 7 | 0.84–0.86 | 0.90–0.93 | 0.66–0.94 | 0.62 / 0.22 |
| no platform factor | 76 | 0 | 0.81–0.82 | 0.63–0.74 | 0.38–0.54 | 0.62 / 0.31 |
| fitted prior | 97 | 7 | 0.85–0.87 | 0.92–0.94 | 0.73–0.97 | 0.62 / 0.20 |
| no platform factor, fitted prior | 77 | 0 | 0.83–0.84 | 0.64–0.76 | 0.40–0.55 | 0.63 / 0.31 |
| 30 markers | 94 | 7 | 0.96–0.97 | 0.81–0.82 | 0.50–0.75 | 0.71 / 0.27 |
| 30 markers, no platform factor<sup>*</sup> | 81 | 0 | 0.85–0.91 | 0.67–0.75 | 0.30–0.49 | 0.70 / 0.33 |
| 30 markers, fitted prior | 95 | 7 | 0.96–0.98 | 0.90–0.94 | 0.55–0.90 | 0.71 / 0.24 |

### Supplementary Methods: datasets

Gut mucosa, Xenium: 16 sections of 11 patients on the 377-gene multi-tissue panel (“337” in the file names), and 4 patients imaged in four configurations, the multi-tissue panel without and with the segmentation stain, the colon panel, and the 5K panel. Masks: nucleus expansion in 20 sections (the 16 sections, whose files predate Xenium Onboard Analysis 2.0, and those imaged without the stain, 5 µm), the stain in 12 (from the segmentation_method column of the cell tables where present). Renal cell carcinoma, Xenium Protein: donor 1, with 28 custom genes (Xenium Onboard Analysis 4.0.0, run May 2025; median 70 transcripts per cell); donor 2, tissue from AcePix Biosciences, with 100 custom kidney genes (Xenium Onboard Analysis 4.0.1, run January 2026; median 54 transcripts per cell), and the vendor’s cell groups (cell_groups.csv). Same chemistry, segmentation stain, and pixel size; only the files needed were extracted from the bundles. Uterine cancer, MERSCOPE (Vizgen FFPE showcase, HumanUterineCancerPatient2-ROCostain): outlines in seven planes; CD45RO signal per cell in cell_metadata.csv; reference, Census uterus (endometrium and myometrium of normal donors; adnexa and decidua excluded). Liver, CosMx, 1000-gene Universal Cell Characterization panel with 14 custom genes, four slides holding ten biopsies and explants: tissue pieces separated by gaps between fields of view (15 pieces of at least 2000 cells; 4 without nuclear counts excluded; fields of view with fewer than a quarter of transcripts in the nucleus removed; the mapping of pieces to patients is not deposited, so pieces are the units); reference, Census liver (normal donors; Census has no steatotic liver disease), 207 over-detected probes excluded as for CosMx lung. In liver, where the platform factor collapsed the labels, the 30 most specific genes per reference type and no platform factor (chosen among seven settings as the one without collapse that agreed best with a reference-free call from canonical markers; Supplementary Note 2).

**Table S20.** Labels lost by every cell class. Kept ratio of each class, whole-cell counts at the depth of nuclear half B against half B, from the stored labels: among cells whose half A label is the class, the share that keep it in the test over the share that keep it in half B, pooled per donor or patient (cells labeled in all three count sets; units with at least 100 cells kept in half B). Median over units, with the range and the number of units; one is no loss. The kept ratio counts labels lost, not labels gained, so classes that absorb others can sit at or above one. Class names follow each tissue’s reference; –, not in the reference or fewer than 100 cells kept in every unit.

| class | gut | kidney | synovium | lung |
|---|---|---|---|---|
| *immune* |  |  |  |  |
| T/NK | 0.71<br>(0.32–0.84; 11) | 0.76<br>(0.60–0.83; 13) | 0.76<br>(0.66–0.86; 16) | 0.57<br>(0.54–0.62; 4) |
| B | 0.91<br>(0.83–0.95; 8) | 0.88<br>(0.75–0.91; 7) | 0.81<br>(0.68–0.89; 10) | 0.77<br>(0.67–0.85; 4) |
| plasma | 0.90<br>(0.79–0.97; 9) | 1.05<br>(0.99–1.12; 9) | 0.96<br>(0.89–1.00; 11) | – |
| mast | 0.84<br>(0.83–0.91; 5) | 0.82<br>(0.71–0.84; 3) | – | 0.79<br>(0.58–0.93; 4) |
| myeloid | 0.96<br>(0.89–1.03; 11) | 0.93<br>(0.84–0.99; 22) | 0.92<br>(0.82–0.96; 17) | 0.85<br>(0.83–0.87; 4) |
| granulocyte | – | – | – | 0.85<br>(0.77–0.94; 2) |
| *stromal and vascular* |  |  |  |  |
| mural or smooth muscle | 0.84<br>(0.81–1.01; 5) | 0.64<br>(0.52–0.72; 9) | 0.92<br>(0.87–0.95; 17) | – |
| endothelial | 0.98<br>(0.91–1.00; 10) | 0.92<br>(0.86–0.98; 21) | 0.94<br>(0.86–0.97; 17) | 1.01<br>(0.95–1.04; 4) |
| fibroblast or stromal | 0.86<br>(0.76–0.98; 12) | 0.96<br>(0.89–1.04; 22) | 1.02<br>(0.97–1.08; 17) | 0.93<br>(0.92–1.00; 4) |
| neural | 0.95<br>(0.91–0.99; 3) | – | – | – |
| *epithelial and parenchymal* |  |  |  |  |
| epithelial | 1.00<br>(1.00–1.06; 15) | – | – | 0.92<br>(0.81–0.99; 4) |
| tumor | – | – | – | 1.00<br>(0.99–1.01; 4) |
| podocyte | – | 0.95<br>(0.92–0.99; 5) | – | – |
| proximal tubule | – | 1.02<br>(0.96–1.09; 17) | – | – |
| loop of Henle | – | 1.02<br>(1.00–1.06; 21) | – | – |
| distal tubule and collecting duct | – | 0.96<br>(0.75–1.00; 19) | – | – |

**Table S21.**
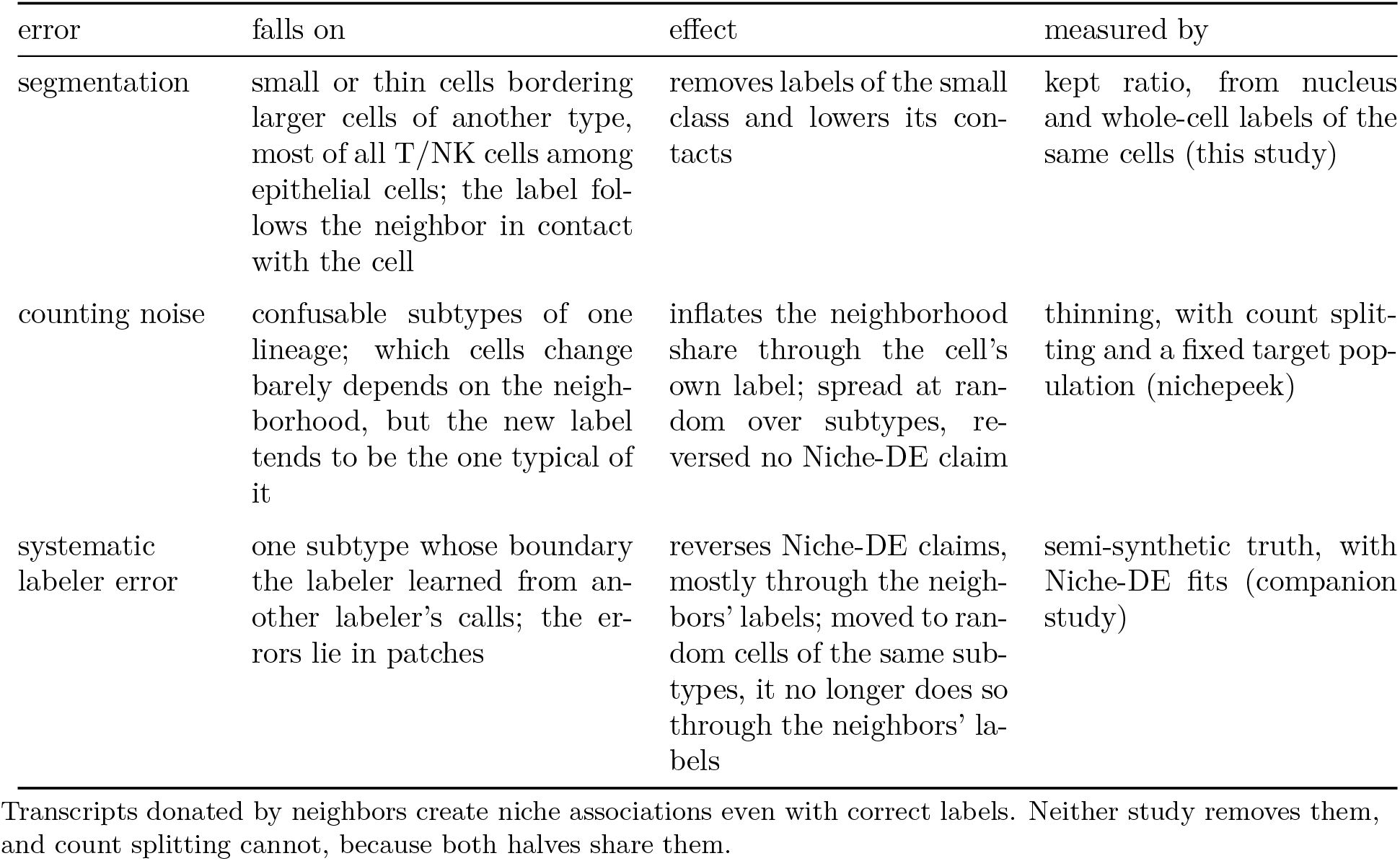
Three kinds of label error. Segmentation error is the error that the kept ratio measures; counting noise and systematic error of the labeler are measured in the companion study [30], by thinning each cell’s counts and by Niche-DE fits on semi-synthetic data. Lung cancer, labeled from whole cells and from nuclei, carries counting noise and segmentation error.

| error | falls on | effect | measured by |
|---|---|---|---|
| segmentation | small or thin cells bordering larger cells of another type, most of all T/NK cells among epithelial cells; the label fol-lows the neighbor in contact with the cell | removes labels of the small class and lowers its con-tacts | kept ratio, from nucleus and whole-cell labels of the same cells (this study) |
| counting noise | confusable subtypes of one lineage; which cells change barely depends on the neigh-borhood, but the new label tends to be the one typical of it | inflates the neighborhood share through the cell’s own label; spread at ran-dom over subtypes, re-versed no Niche-DE claim | thinning, with count split-ting and a fixed target pop-ulation (nichepeek) |
| systematic labeler error | one subtype whose boundary the labeler learned from an-other labeler’s calls; the er-rors lie in patches | reverses Niche-DE claims, mostly through the neigh-bors’ labels; moved to ran-dom cells of the same sub-types, it no longer does so through the neighbors’ la-bels | semi-synthetic truth, with Niche-DE fits (companion study) |

Transcripts donated by neighbors create niche associations even with correct labels. Neither study removes them, and count splitting cannot, because both halves share them.

### Supplementary Methods: label audit

A gene detected in a tissue far above its share in every reference type acts on the classifier as a strong marker of the type with its highest share. Off-target binding of a probe is one cause [54]; a cell state missing from the reference is another. For each section and for the nuclear and whole-cell labels, we listed genes holding at least 0.5% of the counts at more than twice their highest reference share, and counted the calls that exist only through such a gene: calls whose margin over the runner-up is no larger than that gene’s own contribution. For each class we also compared the share of its calls that carry one of its marker transcripts with the share expected of true members at the calls’ depth (support ratio; about one for true members). The audit runs at labeling time; for labels made before it, we recomputed it from the stored labels, runner-ups, and platform factors, which reproduced every stored label. In the first synovium labeling, CCL5 held 0.034–0.139 of nuclear counts, 5–21 times its highest reference share; a median of 0.64 of nuclear T/NK calls existed only through it (up to 0.90), and the T/NK support ratio was 0.41. EEF1G decided 0.02–0.11 of whole-cell B calls. Synovium was relabeled without both genes; in the audited labeling no gene decided more than 0.014 of T/NK calls and the support ratio was 0.92. In the other seven datasets, no over-detected gene decided more than 0.014 of T/NK calls in any section, except TRAC, the T-cell receptor *α* constant gene, in gut (up to 0.19 in one section), and T/NK support ratios were 0.73–1.00. Probe sequences explain one of these genes. The published off-target analysis of the predefined Xenium panels [54] finds all three EEF1G probes of the Prime 5K panel matching EEF1G pseudogenes, and no off-target match for the eight TRAC probes of the gut panels. CCL5 is not among the Prime 5K genes of that analysis, so its probes in the synovium panel were not tested there; the CCL5 probes of five other predefined panels have no off-target match. The cause of the CCL5 excess, therefore, remains open. The dataset authors’ synovium labels, made by clustering within the dataset, agreed with our audited labeling (agreement 0.80) and also kept fewer T cells at contact with large non-immune cells, in 12 of 17 patients, yet their T/NK contacts per 100 fibroblasts barely moved from our nuclear labels (*−*1.8%), whereas our whole-cell labels changed them by *−*23.6%. In kidney, the authors’ labels, made from the same whole-cell counts, call T/NK many cells that our classifier calls myeloid, B, or mast cells.

### Supplementary Methods: extra-change test and masks

Boundary contact used the Xenium cell boundaries of lung, gut, and kidney (a median of 4–6 contacting cells per lung cell). On simulated sections in which every extra change copied the type of a cell 30–100 µm away, never a contacting one, the plain ratio was 1.78–1.89 and the matched ratio 1.00–1.03 (five sections of 60,000 cells each); when changes ignored position, it was 0.98, and when they copied a contacting cell, 3.5 (examples/compartment/regional_sim.py). T/NK nuclei measured 4.9–5.9 µm across (medians per section) and their deposited masks 7.0–11.3 µm; lung tumor nuclei 6.4–9.2 and masks 10.6–18.5 µm. With border shares from polygons, 94–100% of cells per section touch another in lung and gut, 83–98% in kidney, and 97% and 89% in the two renal cell carcinomas.

### Supplementary Methods: contact statistics

Placement: the real number of changes *a → b* beyond the null applied to the half B labels, choosing cells at random, by lowest margin toward *b*, by number of type-*b* contacts, or by a logistic flip model fitted leaving the unit out; scored by the sign of the colocalization change (*k* = 10) for pairs with a real change of at least 0.02. Contacts scored against protein (renal cell carcinomas; protein_contacts.py): one contact graph over all cells with protein (six nearest within 30 µm); partners, cells CD45-negative by protein, or PanCK-positive; T cells called by CD3E protein (positivity as above; and, as a check, the strict T class) or by each labeling (nucleus, whole cell, half B, whole-cell counts at the depth of half B, and the hybrid); contacts per 100 partner cells and the share of T cells contacting a partner, each labeling’s value over the protein value, and whole-cell counts over half B at equal depth, with 95% intervals from the same tile bootstrap applied to both terms of each ratio (200 µm tiles, 1000 draws). Cells that a labeling left unlabeled count as non-T in that labeling; the same ratios on the cells labeled by every count set are in the result tables.

### Supplementary Methods: example cell

For Fig. 1a, the cells of one lung section whose nuclear label is T/NK, whose whole-cell label is tumor and which have a tumor cell among their contact neighbors were ranked by T-cell marker transcripts over the nucleus and tumor-marker transcripts outside it (quality value at least 20); the cell shown ranked first, and the other candidates are listed with the code (claimbreak cell_crop.py).

### Supplementary Methods: neighbor RNA checks

Positional check, lung: before contact was computed, every cell was moved by an isotropic Gaussian error of 5, 10 or 20 µm root mean square, with all cells and with 40% removed at random (the same cells at every level); counts, markers and loads are unchanged. Checks for mislabeled cells, lung and gut: the load was also counted in nuclear half B, rebuilt from the labeling’s genes and seed (its complement, half A, reproduced the recorded half A counts in every section), with whole-cell and nuclear loads then restricted to the label genes; measured T/NK cells were limited to labels with a margin of at least 20 nats, or in addition to at least three nuclear counts of their own markers; the partner cells that define contact were all cells with the partner label. Shape of the distribution: per section, contact over away at the median nuclear load, and the probability that a cell at contact carries a higher load than a cell away (ties counted half). Scripts: examples/compartment/contact_contamination.py (--jitter, --drop-frac, --measure nucB, --min-margin, --min-own, --protein), contact_mislabel.zsh and slidetags_tcr.py contamination.

### Supplementary Methods: transcript-based segmentation

Proseg cells mapped to Xenium nuclei: 70,513 of 73,040 cells in L1 and 175,327 of 189,958 in L3; in the renal cell carcinomas, 443,605 of 465,534 and 636,976 of 690,322. Baysor cells: 64,758 of 73,040 cells in L1, 166,102 of 189,958 in L3, 424,739 of 465,534 and 602,608 of 690,322 in the two renal cell carcinomas.

### Supplementary Methods: protein

Published labels (donor 2): the vendor’s cell groups mapped to T (all T-cell groups), B, and myeloid by name and scored on the cells they cover against the same protein classes and bins, with our labels scored on the same cells (xenium_protein.py groups-border); the groups were made with protein for immune and tumor cells [21], so their recall is not independent of the protein truth. Cell-specificity: in both renal cell carcinomas, CD3E-positive cells carried 10–12 times more T-cell transcripts than other cells, and their nearest neighbors only 2.4–2.6 times more. For each protein, the counts of a class signature (the class’s marker genes present in the panel; for T cells CD3E, CD3D, CD2, TRAC, LCK, CD8A, and others) per 1000 counts were compared between protein-positive and other cells, and between their nearest neighbors (xenium_protein_check.py); the ratio of the two folds was 2.1–4.6 for CD3E, CD8A, CD45, and CD68 in both sections. Strict protein classes (xenium_protein.py truth --strict): T, CD3E^+^ CD45^+^ cells without CD68 or pan-cytokeratin; B, CD20^+^ CD45^+^ cells without CD3E, CD68, or pan-cytokeratin; positive calls at posterior 0.5 and 0.9. With these classes the T recall ratio was 0.92 (compared with 0.91) in donor 1 and 0.97 (compared with 0.96) in donor 2.

MERSCOPE nuclei: StarDist (2D_versatile_fluo) on the middle of seven DAPI planes, aver-aged in 2*×*2 blocks (0.216 µm per pixel), in 442-µm tiles with overlap. Per tile, contrast from the 1st and 99.8th percentiles of imaged pixels (unimaged areas are zero), then contrast-limited adaptive histogram equalization (14-µm window, clip limit 0.01) and Gaussian smoothing (0.2 µm); objects under 15 µm^2^ dropped. The settings were chosen on six 324-µm test crops (dense, bright, hazy, sparse, and partly unimaged tissue) before the section was segmented once (merscope_crops.py, merscope_nuclei.py, merscope_owners.py). 615,062 nuclei. Of the transcripts in cells, 0.48 were nuclear, and 0.08 lay in another cell’s nucleus and count as cytoplasm (merscope_to_h5ad.py). CD45RO: Ashman *D* = 2.19; 0.10 of cells positive; the raw signal did not separate. Cell-specificity: each stain-high cell against its nearest stain-low neighbor within 15 µm (fold of the T-cell signature), for CD45RO, three membrane stains, poly(A), and DAPI, and for CD45RO on both Xenium Protein sections (merscope_protein_check.py, xenium_protein_check.py); a CD45RO-high cell carried 4.9 times more T-cell transcripts than its nearest CD45RO-low neighbor; the comparison with all nearest neighbors used for Xenium Protein above understates cell-specificity where T cells lie in aggregates (1.62 here). Labels as for the Xenium sections; a second labeling excluded the 18 genes that the label audit flagged as over-detected (PKM, IDH1, AKT1, HDAC1, ERBB2, and others; relabel_excluding.py).

### Supplementary Methods: labels against protein

Sizes are reported as the kept ratio throughout; the share of T/NK labels among all labels (wholeB over nucB) ran below protein along the expansion series and is used only for the pre-specified synovium outcome E3. Overlap control: on cells labeled in A, B, and the whole cell, the kept ratio split by protein status and the recall ratio split by the half A label, with the positivity calls of the protein runs. When the two ratios agree: write the kept ratio as *K* = [*πk_w_*(*T*) + (1 *− π*)*k_w_*(*F*)]*/*[*πk_n_*(*T*) +(1 *− π*)*k_n_*(*F*)], with *π* the share of true class members among the half A calls of the class and *k_w_*, *k_n_* the shares of true (*T*) and false (*F*) calls that the whole-cell counts and half B keep. Half A is independent of half B and of the rest of the cell, so *K* equals the protein marker recall ratio when false calls are rare or are kept at the same relative rate as true ones. False calls that both nuclear halves repeat, such as those of an over-detected probe, raise *k_n_*(*F*) more than *k_w_*(*F*), and *K* falls below the protein ratio. Ground-truth simulation (kept_ratio_sim.py): sections of 30,000 cells (T cells small and RNA-poor, tumor cells large and RNA-rich) with a planted share *ψ* = 0–0.6 of each cell’s cytoplasmic counts (*ψ/*4 of its nuclear counts) drawn from its contacting cells, labeled as the real sections; with and without a probe that reads as a T marker in the reference and holds 15% of fibroblast nuclear counts; five seeds. The true recall ratio reached 0.58; with the probe, the kept ratio was 0.93 with no planted contamination. Without the probe, the kept ratio recovered the true loss to within 0.006. Counting unlabeled cells as missed changes the nuclear T recall to 0.51 and 0.45 (from 0.64 and 0.63; 0.79 and 0.71 of T cells defined by protein markers labeled from the nucleus) and the whole-cell recall to 0.54 and 0.51. The synovium outcomes, including E4 and T3, were written down before the synovium sections were labeled; Supplementary Table S3 gives them for both labelings, each computed from its own result folders. T3 uses the split typing of every foreign-share value; our first evaluation used the rule without it, and the table gives both. Foreign shares and T3 leave out the genes a labeling excluded. Both compare foreign with own marker counts, so they depend on how many own markers the panel holds for a type, and they are compared here only within a panel.

### Supplementary Methods: T cells among epithelial cells by condition

Gut, the 16 sections on the 377-gene panel (one panel and mask type for every patient); conditions from the GEO sample titles (site, disease, inflamed or not; GSE312420), assigned per section. For each labeling (nuclear, whole cell, hybrid) and section: epithelial cells, T/NK cells, and T cells among epithelial cells, T/NK cells with at least half epithelial cells among the labeled cells within 25 µm, all from that labeling; per patient, sections pooled within each condition level (a patient with sections in two levels counts in both). Readouts: T cells among epithelial cells per 100 epithelial cells, and their share of all T/NK cells. Contrasts: ratio of the medians over patients between levels, with 95% intervals from 2000 bootstrap draws of patients within each level, the same draws for every labeling, and the ratio of that ratio between labelings; also within each site and each disease (claimbreak ibd_condition.py). Comparison with histology (ibd_histology.py): per patient, T cells among epithelial cells per 100 epithelial cells from each labeling, pooled over the patient’s sections, and the ratio of the whole-cell and hybrid values to the nuclear value; medians and ranges over patients. Pathologists count lymphocytes within the epithelial layer on CD3-stained sections per 100 surface or villus epithelial cells: a mean of 3.8 in normal terminal ileum [27] and a median of 3.6 in normal colon [26]. Ours are per 100 epithelial cells of every kind, crypts included, in two-dimensional sections and with the inside rule above, and nuclear labels leave cells with too few nuclear counts unlabeled, which lowers their count; IEL counts in IBD also vary with site and activity. These biases run in opposite directions, so agreement in level may be partly coincidental; the comparison that does not depend on histology is the ratio between labelings in each patient. Medians over patients: nuclear 3.53 (0.55–6.53), whole cell 1.36 (0.31–3.79), hybrid 2.97 (0.57–5.58); whole cell over nuclear 0.50 (0.25–0.83), hybrid over nuclear 0.89 (0.84–1.02).

### Supplementary Methods: a contrast between conditions

For a class that mostly loses labels, whole-cell counts at the depth of half B give the count from half B times the kept ratio *K*, so a contrast between groups 1 and 2 is biased by *K*_1_*/K*_2_. In gut, T cells among epithelial cells kept 0.39 of their labels in inflamed and 0.41 in non-inflamed sections, pooled over patients (all T/NK cells, 0.59 and 0.70; ibd_bias.py). Whole-cell counts gained no more T/NK labels than half B did: of the T/NK labels of cells labeled by half A, 0.108 had another half A label, against 0.125 for half B (ibd_decomp.py, gain rows). To see where a labeling changes the inflamed / non-inflamed contrast of T cells among epithelial cells per 100 epithelial cells, we wrote its whole-over-nucleus factor, on counts pooled per condition level, as the ratio of the counts among epithelial cells over the ratio of epithelial counts, and the former as the ratio of T/NK counts times the ratio of the shares of T/NK cells called inside; each term is an inflamed-over-non-inflamed ratio, with a bootstrap over patients within levels (ibd_decomp.py). A T/NK cell was called inside when at least half of the labeled cells within 25 µm were epithelial, by the same labeling. At full depth, the factor was 0.82 (0.64–1.11): the epithelial term was 0.98, and the share of T/NK cells called inside 0.75 (0.59–0.93); as the median over patients, the factor was 0.81. At equal depth, it was 0.91 (0.67–1.15), with a T/NK count term of 0.89 (0.71–1.07), consistent with the ratio of the kept ratios. Leaving out any one section kept the pooled factors at 0.75–0.92 (full depth) and 0.87–0.97 (equal depth). Counting all cells around a T/NK cell instead, with unlabeled cells as not epithelial, mixes how many cells a labeling labels into the region call; with that rule, the full-depth factor was 0.65 (0.46–0.89) pooled and 0.52 as the median over patients. With 11 patients, the contrast’s change with the labeling is not resolved under the labeled-cell rule.

### Supplementary Methods: identity and state of the calls by region

Identity of the calls, per bin and for the partner cells: nuclear and whole-cell counts, the half A margin and runner-up label, and from the cell tables cell and nucleus areas and the nuclear counts of T-cell genes (CD3E, CD3D, CD3G, CD2, CD247, TRAC, TRBC1, TRBC2, CD7, LCK, CD8A, CD8B, IL7R), partner genes, and cytotoxic genes (GZMA, GZMB, GZMH, GZMK, PRF1, NKG7, GNLY) per 1000 nuclear counts, as available in each panel; medians per unit, then over units. State: a call is positive for a gene set (cytotoxic; proliferation; naive-like; exhaustion; activation; CD8; CD4; *γδ*; tissue-resident) with at least one nuclear count of it; the loss of positive and negative calls compared within five strata of equal size of nuclear counts per unit, with Mantel–Haenszel weights *n*_+_*n_−_/*(*n*_+_ + *n_−_*) (strata with at least five calls on each side; units with at least ten on each side), and the same matching for inside against away (region_check.zsh, state_depth.zsh). B and plasma cells as for T/NK cells.

### Supplementary Methods: resolVI

Windows were 2071 and 2394 µm wide, the three per renal cell carcinoma section with the most cells labeled T/NK (120,619 and 137,307 cells). The unsupervised model was trained for 50 epochs, with package defaults otherwise, and the graph had 100 nearest neighbors. The filter kept 0.05–0.38 of the labels per window. As sensitivity analyses, we repeated the relabeling with the authors’ threshold for fine labels (40%, which kept 0.34–0.79 of the labels) and, at 5%, with a second seed. The semi-supervised model drew 30 posterior samples (one seed, one Quadro P4000 GPU).

### Supplementary Methods: synovium reference

From the four RA samples of GSE200815: cells with at least 500 UMIs, 250 genes, and a mitochondrial share below 0.25; 2000 variable genes, 30 principal components, Harmony by sample [55], Leiden clustering [56]; clusters assigned to coarse classes by lineage markers and reviewed by hand; a mixed B and plasma cluster split by lineage score; at most 3000 cells per class. The same pipeline without Harmony gave the same results except for B cells. The synovium sections were labeled without CCL5 and EEF1G, the two genes the label audit found deciding calls in the first labeling (Methods); every other setting was kept (claimbreak relabel_excluding.py).

